# A glial circadian gene expression atlas reveals cell type and disease-specific reprogramming in response to amyloid pathology or aging

**DOI:** 10.1101/2024.05.28.596297

**Authors:** Patrick W. Sheehan, Stuart Fass, Darshan Sapkota, Sylvia Kang, Henry C. Hollis, Jennifer H. Lawrence, Ron C. Anafi, Joseph D. Dougherty, Jon D. Fryer, Erik S. Musiek

**Affiliations:** Department of Neurology, Washington University School of Medicine, Saint Louis MO, USA; Department of Genetics, Washington University School of Medicine, Saint Louis MO, USA; Department of Psychiatry, Washington University School of Medicine, Saint Louis MO, USA; Department of Biological Sciences and Department of Neuroscience, University of Texas at Dallas, Richardson, TX, USA; Department of Neuroscience, Mayo Clinic, Jacksonville, FL, USA; School of Biomedical Engineering and Health Systems, Drexel University, Philadelphia, PA, USA; Department of Medicine, Chronobiology, and Sleep Institute, Perelman School of Medicine, University of Pennsylvania, Philadelphia, PA, USA; Intellectual and Developmental Disabilities Research Center, Washington University School of Medicine, St. Louis, MO, USA; Department of Neuroscience, Mayo Clinic, Scottsdale, AZ, USA; Center on Biological Rhythms and Sleep, Washington University School of Medicine, St. Louis, MO, USA

**Keywords:** circadian, transcriptome, glia, astrocyte, microglia, Alzheimer’s Disease, aging

## Abstract

While circadian rhythm disruption may promote neurodegenerative disease, how aging and neurodegenerative pathology impact circadian gene expression patterns in different brain cell types is unknown. Here, we used translating ribosome affinity purification methods to define the circadian translatomes of astrocytes, microglia, and bulk cerebral cortex, in healthy mouse brain and in the settings of amyloid-beta plaque pathology or aging. Our data reveal that glial circadian translatomes are highly cell type-specific and exhibit profound, context-dependent reprogramming of rhythmic transcripts in response to amyloid pathology or aging. Transcripts involved in glial activation, immunometabolism, and proteostasis, as well as nearly half of all Alzheimer Disease (AD)-associated risk genes, displayed circadian oscillations, many of which were altered by pathology. Amyloid-related differential gene expression was also dependent on time of day. Thus, circadian rhythms in gene expression are cell- and context dependent and provide important insights into glial gene regulation in health, AD, and aging.

## Introduction

Circadian rhythms are 24-hour oscillations in behavior, physiology, and gene expression which are entrainable, persist under constant conditions, and are driven by core circadian clock proteins. Numerous studies have demonstrated robust circadian rhythms in gene expression which involve not just core clock proteins, but thousands of other genes which are highly tissue-specific^1, 2^. Several reports suggest that the population of transcripts exhibiting circadian oscillation may change substantially in the presence of different disease states or insults, a phenomenon known as circadian reprogramming^3^. This shift in rhythmic gene expression has been described in multiple organs and disease states, including mouse hippocampus in epilepsy^4^, and drosophila and human brain tissue in aging^5, 6^. However, most circadian transcriptomic studies have relied on bulk homogenized tissue, and only a few have attempted to dissect unique circadian transcriptomes of specific cell types, particularly in a complex, heterogeneous tissue such as the brain^7, 8^. Thus, little is known about cell type specific circadian gene expression in the brain, either in healthy or diseased tissue.

Aging and neurodegenerative diseases are both associated with changes in circadian rhythms. Behavioral rhythms become fragmented with aging in mice and humans, in parallel with blunting of circadian electrical output from the suprachiasmatic nucleus of the hypothalamus, the central clock of the body^9, 10^. Circadian function is further disrupted in age-related neurodegenerative diseases such as Alzheimer’s Disease (AD), which is characterized by decreased circadian amplitude and fragmentation of behavioral rhythms^11, 12^. Circadian rhythm fragmentation begins to occur prior to the onset of AD symptoms, and can be seen in cognitively-normal people with biomarker evidence of amyloid plaque pathology^13, 14^. Glia cells in particular exhibit robust cell-autonomous circadian rhythms, and genetic disruption of glial circadian clocks can influence disease-relevant processes such as glial activation, endolysosomal function, inflammation, and protein aggregation^17–24^. While most studies focus on behavioral circadian rhythms, little is known about how aging or early AD pathology, particularly amyloid plaques, may influence circadian gene expression in different cell types in the brain, or if this process might contribute to AD pathogenesis.

Here, we have used translating ribosome affinity purification (TRAP) methods to elucidate cell-specific circadian translatomes of astrocytes and microglia, as well as bulk cortex, at high temporal resolution under constant conditions. We have then examined the impact of amyloid plaque pathology or aging on these cell-specific circadian gene expression patterns. We observed that astrocytes and microglia have unique, cell type-specific circadian translatomes in vivo which change dramatically and uniquely in the setting of either amyloid plaques or aging. Amyloid pathology alters circadian rhythmicity of microglial homeostatic and diseases-associated activation markers, while numerous AD-related transcripts gain rhythmicity in astrocytes. Aging induces striking rhythms in endolysosomal pathways and mTOR expression in astrocytes, but blunts core clock oscillations in microglia. Finally, we find that time of day of harvest influences the analysis of differential gene expression in response to amyloid pathology. We provide this rich dataset as a publicly-available, fully-searchable website that should provide unique insights into the role of circadian rhythms in brain health and disease, and inform future transcriptomics efforts in AD.

## Results

### Assessment of TRAP-RNAseq for defining glial circadian translatomes in vivo

To assess cell-specific circadian rhythms in gene expression in vivo, we utilized astrocyte-specific TRAP mice (*Aldh1l1*-RPL10a^eGFP^, termed AstroTRAP) and microglia-specific RiboTag mice (*Cx3cr1*-Cre^ERT2^;LSL-Rpl22^HA^; termed mgRiboTag) to isolated cell-specific ribosomes (and associated RNA) for transcriptomics^25, 26^ (Fig 1a). The mgRiboTag mouse received tamoxifen 60 days prior to harvest to allow for repopulation of peripheral macrophages and increased cell-specificity of recombination in microglia^27^.We entrained mice to 12h:12h light:dark cycles, then placed them in constant darkness for 24 hours prior to and during the experiment. We sacrificed mice in the dark at two hour intervals over a 24 hour period in duplicate, perfusing with cycloheximide to halt ribosomal activity and capture ribosome-associated mRNA levels exactly at the time of perfusion. Of note, we repeated the experiment months later and combined the data from both studies after batch correction, demonstrating reproducibility of the study (Fig. S1A). A sample of cerebral cortex lysate that was not subjected to immunoprecipitation (Pre-IP) was also sequenced as the bulk cortex control. Utilizing this approach, we compared cell type-specific transcript levels between Pre-IP, AstroTRAP, and mgRiboTag samples, and found enrichment of astrocyte-specific transcripts by 5-10 fold in the AstroTRAP mice and microglia-specific transcripts by 10-20 fold in mgRiboTag mice (Fig 1C, F). In order to assess the impact of amyloid plaque pathology on the circadian “translatomes” of astrocytes and microglia in vivo, we crossed AstroTRAP and mgRibotag mice to APP/PS1-21 mice, which begin to develop amyloid plaque pathology starting at 2 months of age ^28^. These mice, termed AstroTRAP-APP and mgRiboTag-APP, were harvested alongside their AstroTRAP and mgRiboTag littermates at six months of age, when there is robust amyloid plaque deposition in the cortex (Fig 1E). The TRAP/RiboTag enrichment for cell-specific transcripts was similar in WT and APP/PS1 mice (Fig 1D, G). These results demonstrate the suitability of our system to identify circadian patterns in astrocyte- and microglia-specific gene expression in vivo.

**Fig. 1:**
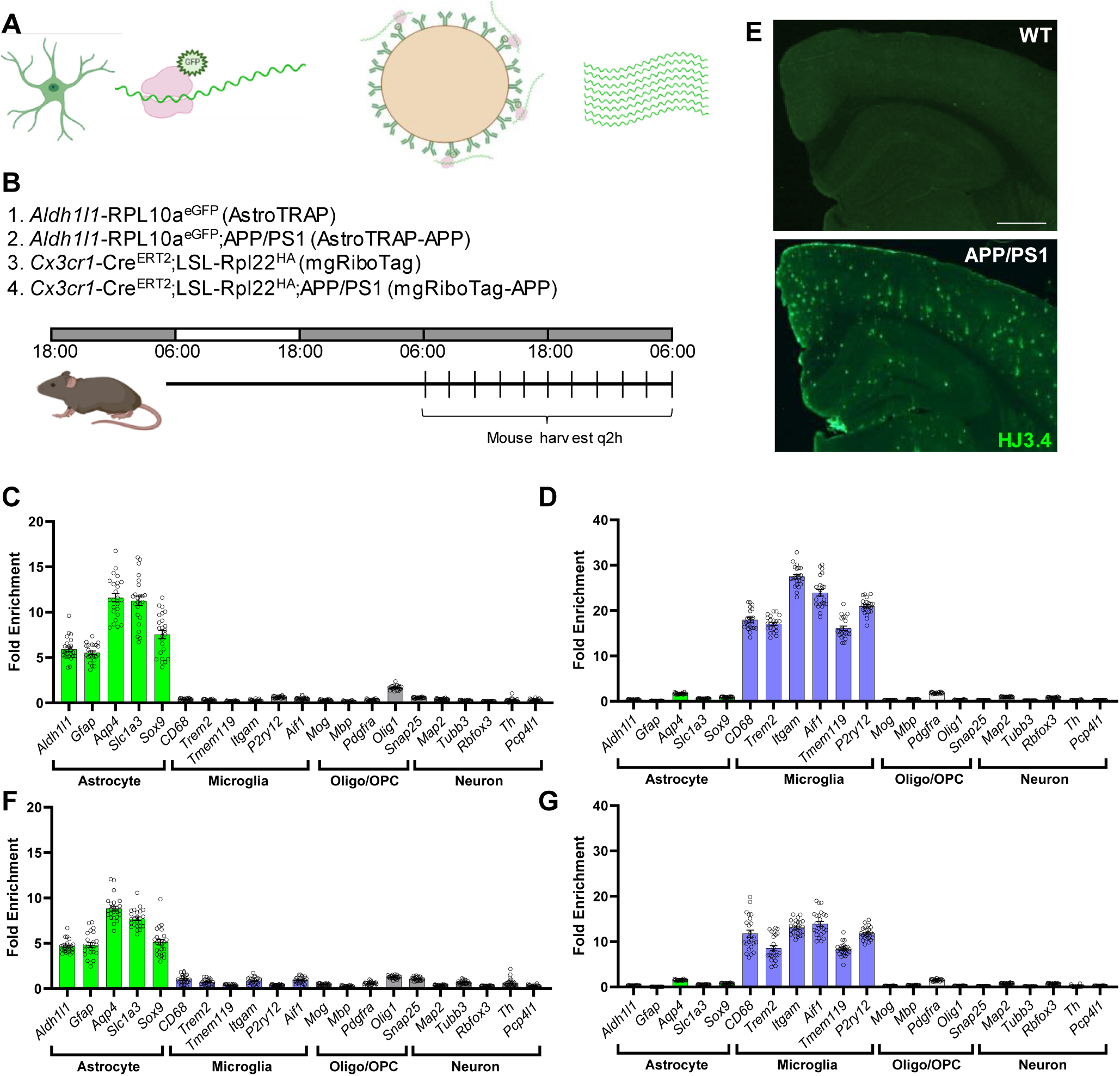
Analysis of circadian rhythms in astrocytes and microglia using Translating Ribosome Affinity Purification (TRAP)/RiboTag. A. Schematic showing the steps in TRAP/RiboTag. B. Listing of the mouse lines used and schematic of the lighting paradigm and mouse harvest schedule. C,D. Fold enrichment of cell type specific gene expression (TRAP/Pre -IP) from AstroTRAP mice (C) and mgRiboTag mice (D). E. Representative image of amyloid plaque burden in a 6mo APP/PS1 - 21 mouse, as assessed byanti-Aβ antibody HJ3.4b. F,G Fold enrichment of cell type specific gene expression (TRAP/Pre -IP) from AstroTRAP-APP mice (F) and mgRiboTag-APP mice (G).

### Amyloid pathology causes circadian reprogramming in bulk cerebral cortex with preservation of the core clock

We first analyzed our Pre-IP samples to assess circadian rhythms in gene expression in bulk cerebral cortex of WT and APP/PS1-21 mice. We used CPM values to analyze rhythmicity using Rhythmicity Analysis Incorporating Nonparametric methods (RAIN), a commonly used algorithm for identifying circadian oscillation in transcript expression in RNAseq data ^29^. Due to the high temporal resolution of our dataset, we used a RAIN adjusted P value <0.01 to identify rhythmically expressed genes. We plotted rhythmic genes in WT and APP/PS1 bulk cortical tissue and organized them based on acrophase (time of peak expression). These plots clearly demonstrate that many transcripts which were identified as rhythmic in WT cortex did not appear rhythmic in APP/PS1 brain, while a subset of transcripts which were not rhythmic in WT mice did gain rhythmicity in APP/PS1 cortex (Fig. 2A). We next used KEGG pathway analysis to identify pathways which show circadian regulation in bulk cortex, using the list of rhythmic genes identified by the RAIN analysis above. When examining transcripts which were rhythmic in both WT and APP/PS1 tissue, we found enrichment for pathways related to MAPK signaling, PI3K signaling, lipid/atherosclerosis, and circadian rhythms (Fig. 2B). Transcripts which were rhythmic only in WT tissue were enriched for lysosome and autophagy genes (Fig 2C), suggesting that amyloid disrupts normal circadian regulation of protein degradation pathways. Transcripts which were rhythmically expressed in APP/PS1 cortex were enriched in hormone synthesis pathways and inflammatory NF-kappaB signaling (Fig 2D). A full list of rhythmic KEGG pathways in all datasets is available in the Supplemental Table 1. The rhythmicity of specific pathways suggests that there may be coordinated expression across genes in order to produce a rhythmically-controlled process. To test this, we plotted all genes that were identified as rhythmic in WT cortical tissue in the “Lysosome” KEGG pathway in a heatmap. Interestingly, the majority of rhythmic lysosome genes peaked at the same time of the day, suggesting that this process may be rhythmically controlled due to the coordinated rhythmic expression of lysosome genes (Fig 2E). The other rhythmic pathways in WT and APP/PS1 tissue showed similar coordination in phase of rhythmic gene expression (Fig S1B,C) Finally, we plotted the phase of expression of each rhythmic gene in WT and APP/PS1 tissue and found that rhythmic genes in WT bulk cortical tissue peaked in a biphasic pattern, while APP/PS1 rhythmic genes largely peaked at one time of day (CT10) (Fig. 2F).

**Fig. 2:**
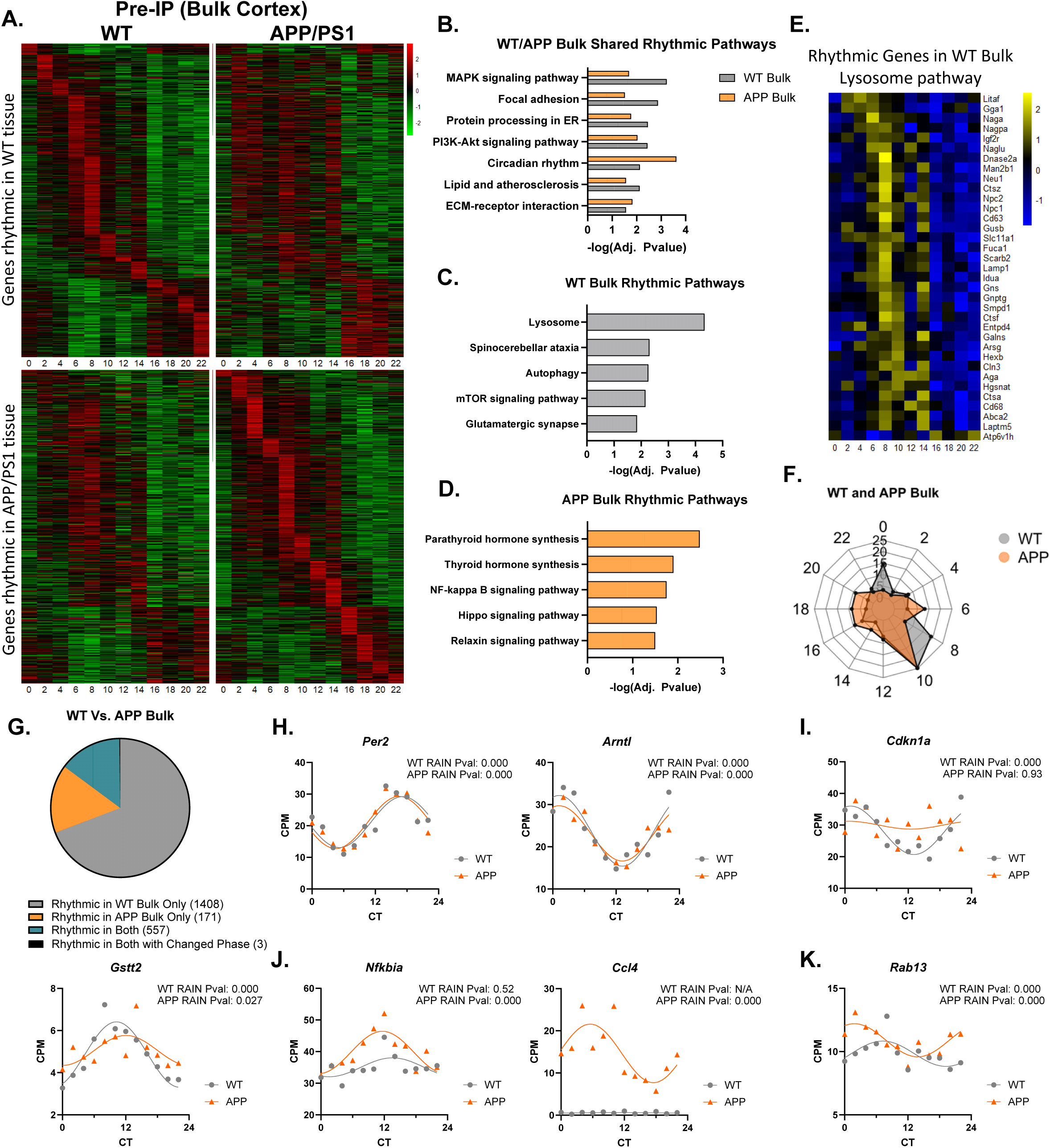
Circadian transcriptional rhythms and reprogramming of bulk cortex transcripts in WT and APP/PS1 mice. A. Heatmaps showing transcripts that were rhythmic in bulk cortex tissue from WT mice (upper) and rhythmic in APP/PS1 mice (lower). In both rows of heatmaps, the genes plotted are in the same order to compare differences in rhythmic expression between mice. B-D. KEGG pathway analysis of cortex transcripts identified as rhythmic (by RAIN analysis) in (B) both WT and APP/PS1 mice, (C) WT mice, or (D) APP/PS1 mice. E. Heatmap showing temporally-coordinated expression of KEGG lysosome pathway genes in bulk cortex from WT mice. F. Radar plot showing acrophase distributions of rhythmic bulk cortex transcripts from WT (grey) or APP/PS1 (orange) mice. G. Pie chart depicting the number of transcripts that gained or lost rhythms in different datasets, based on compareRhythms analysis of rhythmic transcripts in bulk tissue. A total of 2139 were identified as rhythmic by RAIN analysis across all datasets. H-K. Graphs showing circadian expression patterns of transcripts from bulk cortex from WT (grey) or APP/PS1 (orange) mice. H. Core clock genes *Per2* and *Arntl* (Bmal1) remained rhythmic. I. Senescence marker *Cdkn1a* and glutathione transferase *Gstt2* lost rhythmicity in APP/PS1. J. Inflammatory transcripts *Nfkbia* and *Ccl4* gained rhythmicity in APP/PS1. K. Endosomal gene *Rab13* changed phase in APP/PS1. Adjusted P values from RAIN are shown. Each datapoint is the average of two mice, each from separate experiments.

To compare whether genes have gained or lost rhythmic expression in WT and APP/PS1 cortical tissue in a more statistically rigorous manner, we analyzed all transcripts which were identified as rhythmic by RAIN using *compareRhythms*, an R package used to identify differential circadian rhythmicity between conditions (Fig 2G) ^30^. First, we identified 557 genes that were consistently rhythmic in both datasets. Among these were the core circadian clock genes, such as *Arntl*, *Per2*, *Ciart*, and *Nr1d1* (Fig 2H, S1D), which appeared to be unperturbed by amyloid pathology. These data also illustrate the robust nature of the dataset, as all core clock genes show appropriate phase and strong rhythmicity, as compared to other datasets^1^. Next, we identified 1408 transcripts which were rhythmic specifically in WT tissue but lost rhythms in APP/PS1 tissue. All of these genes, such as the cellular senescence marker *Cdkn1a*, the glutathione-s-transferase *Gstt2*, and synaptic proteins *Homer1* and *Synj2*, showed loss or blunting of circadian expression rhythms in APP/PS1 tissue (Fig 2I and S1E). Finally, 171 transcripts gained rhythmicity in APP/PS1 tissue, as they were rhythmic only in the presence of amyloid plaques and were not normally rhythmic in WT tissue. These included inflammatory mediators such as the NF-kappaB component *Nfkbia* and chemokine *Ccl4* (Fig 2J). Interestingly, there were only 3 genes that were rhythmic in both datasets and changed phase of expression, such as *Rab13*, a gene involved in vesicular transport (Fig 2K). These data show that while core clock gene oscillation is preserved, the circadian transcriptome of the cerebral cortex is perturbed by amyloid pathology, with the majority of transcripts losing rhythmicity, while a smaller group undergoes circadian reprogramming to gain rhythms in the setting of β-amyloidosis. In general, genes that lost rhythm were involved in autophagy and lysosomal function, while genes gaining rhythmicity were inflammatory.

### The astrocyte circadian translatome is reprogrammed in response to amyloid pathology to mediate rhythmic expression of AD-related transcripts

We next performed a similar analysis of rhythmic gene expression on astrocyte-enriched, ribosome-associated transcripts from the AstroTRAP and AstroTRAP-APP mice. Again, many transcripts were rhythmic in astrocytes from WT but did not appear rhythmic in APP/PS1 mice, while others gained rhythms in the setting of amyloid plaques (Fig 3A). KEGG pathway analysis of all rhythmic transcripts in both WT and APP/PS1 astrocytes identified the core circadian clock as preserved in both (Fig. 3B). Transcripts involved in metabolism, glutathione metabolism, and insulin signaling were rhythmic in astrocytes of WT mice (Fig. 3C), but not in astrocytes of APP/PS1 mice. Meanwhile, FoxO, Notch, and PI3K signaling pathways, along with inflammatory TNF signaling, were identified as rhythmic exclusively in APP/PS1 tissue (Fig. 3D). Once again, we identified temporally-coordinated gene expression across genes within these rhythmic pathways, exemplified by the insulin signaling pathway in WT Astrocytes (Fig 3E, S2A,B). Finally, we plotted the phase of circadian gene expression in both WT and APP/PS1 astrocytes and found that astrocytes in WT tissue were biphasic in their circadian gene expression, while astrocytes in APP/PS1 tissue lost this coordinated gene expression (Fig 3F).

**Fig. 3:**
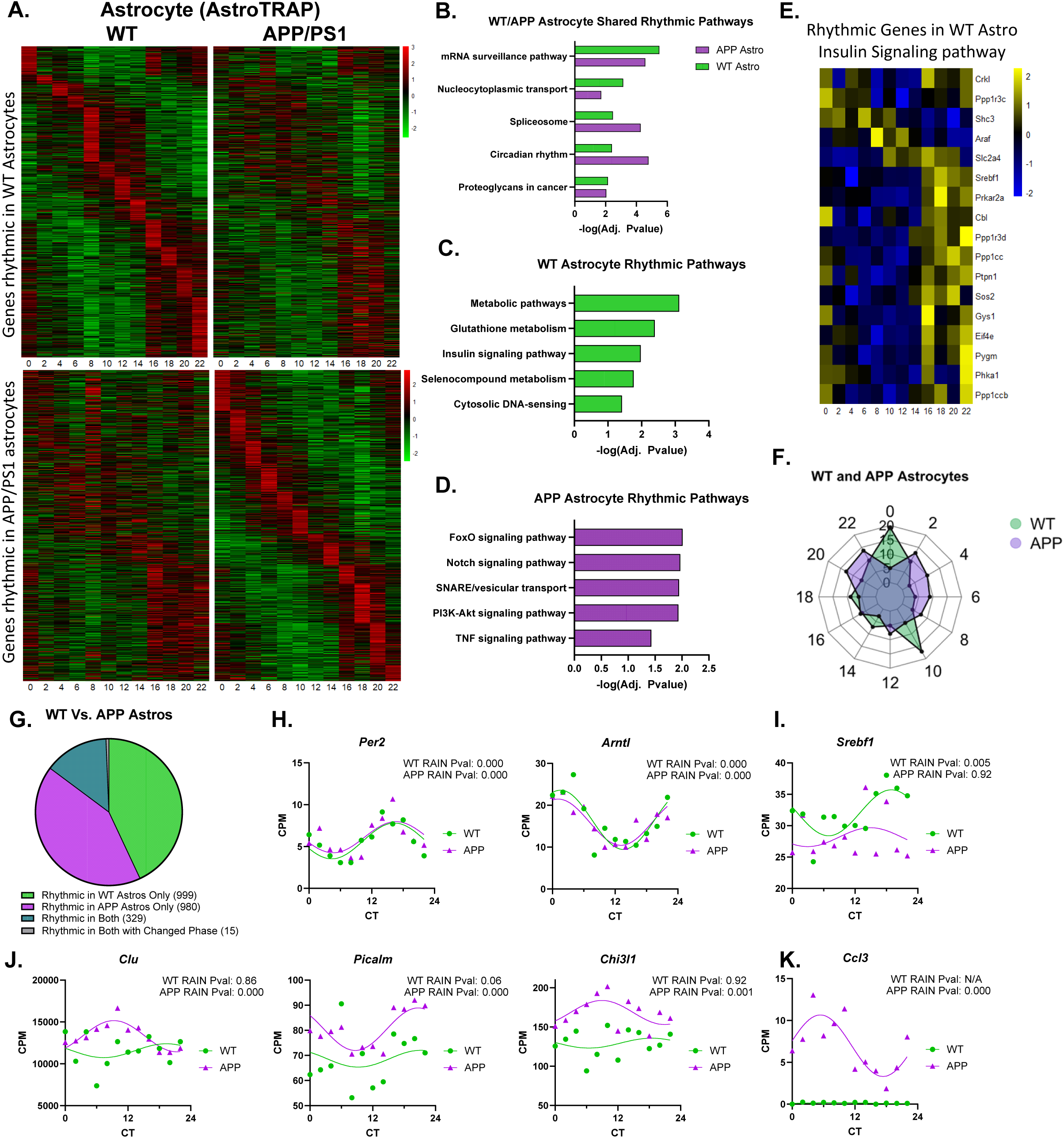
Astrocyte circadian translatomes and reprogramming in WT and APP/PS1 mice. A. Heatmaps showing transcripts that were rhythmic in WT astrocytes (upper) and rhythmic in astrocytes from APP/PS1 mice (lower). In both rows of heatmaps, the genes plotted are in the same order to compare differences in rhythmic expression between mice. B-D. KEGG pathway analysis of astrocyte transcripts identified as rhythmic (by RAIN analysis) in (B) both WT and APP/PS1 mice, (C) WT mice, or (D) APP/PS1 mice. E. Heatmap showing temporally-coordinated expression of KEGG insulin signaling pathway genes in astrocytes from WT mice. F. Radar plot showing acrophase distributions of rhythmic astrocyte transcripts from WT (green) or APP/PS1 (purple) mice. G. Pie chart depicting the number of transcripts that gained or lost rhythms in different datasets, based on compareRhythms analysis of rhythmic transcripts in microglia. Total of 2323 were identified as rhythmic by RAIN analysis across all datasets. H-K. Graphs showing circadian expression patterns of transcripts from microglia from WT (green) or APP/PS1 (purple) mice. H. Core clock genes Per2 and Arntl (Bmal1) remained rhythmic. I. Cholesterol response gene Srebf1 lost rhythmicity in APP/PS1. J & K. AD-related transcripts Clu, Picalm, and Chi3l1, as well as chemokine Ccl3 (K), gained rhythmicity in APP/PS1. Adjusted P values from RAIN are shown. Each datapoint is the average of two mice, each from separate experiments.

We next performed *compareRhythms* analysis and found that, unlike the bulk cortex dataset, astrocytes did not have reduced rhythmic gene expression in APP/PS1 mice (Fig. 3G). Instead, 999 genes that were specifically rhythmic in WT astrocytes lost rhythms in APP/PS1, while a similar number of genes (980) that were not rhythmic in WT mice gained rhythms in APP/PS1 mice. It is notable that most of these rhythmic transcripts were not identified in bulk cortex tissue, illustrating the need to look at specific cell types. Only 329 transcripts were rhythmic in both WT and APP/PS1 astrocytes, and this group once again included the core circadian clock genes, showing that the core clock is robust in astrocytes in the presence of amyloid plaque pathology (Fig 3H and S2C). We identified genes such as *Srebf1*, lipid synthesis regulator, which was rhythmic in WT astrocytes and lost rhythms in APP/PS1 astrocytes (Fig. 3I). To our surprise, a number of AD-associated transcripts, including several GWAS-identified AD risk factors, gained rhythms in astrocytes in APP/PS1 mice. For example, *Clu* (encoding Clusterin/ApoJ), *Picalm*, and *Chi3l1* (encoding inflammatory biomarker YKL-40) all gained prominent rhythms in APP/PS1 astrocytes (Fig 3J). Other genes were not expressed in WT astrocytes, but were upregulated, and gained rhythmic expression in APP/PS1 astrocytes, such as the chemokine *Ccl3*, which is dysregulated in AD (Fig 3K) ^31^. These data show that astrocytes have hundreds of rhythmically expressed genes, many of which are perturbed by amyloid plaques. The core clock of astrocytes is robust in the setting of amyloid pathology, but clock-controlled transcripts are reprogrammed, with several prominent AD-related transcripts gaining rhythmicity in the setting of amyloid.

### The microglial circadian translatome includes many canonical activation and homeostatic genes and is disrupted by amyloid plaques

We next examined circadian rhythms in microglia-enriched ribosome-associated RNAs, using mgRiboTag and mgRiboTag-APP mice and similar methods as above. As in the other datasets, many microglia genes appeared to lose or gain rhythms in the setting of amyloid plaque pathology (Fig. 4A). KEGG pathways analysis of rhythmic transcripts showed a number of critical signaling pathways that were rhythmic in microglia in both WT and APP/PS1 mice, including mTOR signaling, TNF signaling, HIF-1α and VEGF signaling (Fig. 4B). Metabolic pathways were the most prominent rhythmic pathways in all microglia. Transcripts which were rhythmic specifically in microglia of WT mice included Huntington, Parkinson, Alzheimer, and prion disease pathways, spinocerebellar ataxia, and ALS (Fig. 4C). Lysosome and proteasome pathways were also rhythmic in WT microglia. However, the rhythmicity of these pathways was lost in microglia from APP/PS1 mice, while pathways involved in PI3K-Akt signaling, lipids, and ferroptosis became rhythmic (Fig 4D). We plotted rhythmic genes in WT microglia involved in multiple pathways (Fig. S3A,B), including proteasome (Fig. 4E), Alzheimer’s disease (Fig. S3C), oxidative phosphorylation (Fig. S3D), and reactive oxygen species (Fig. S3E), and found that transcripts within each pathway exhibited temporally-coordinated expression. When examining gene expression phase, we found that microglial transcripts, unlike astrocytes, had a broad distribution of acrophases, and this did not change in the setting of amyloid pathology (Fig. 4F).

**Fig. 4:**
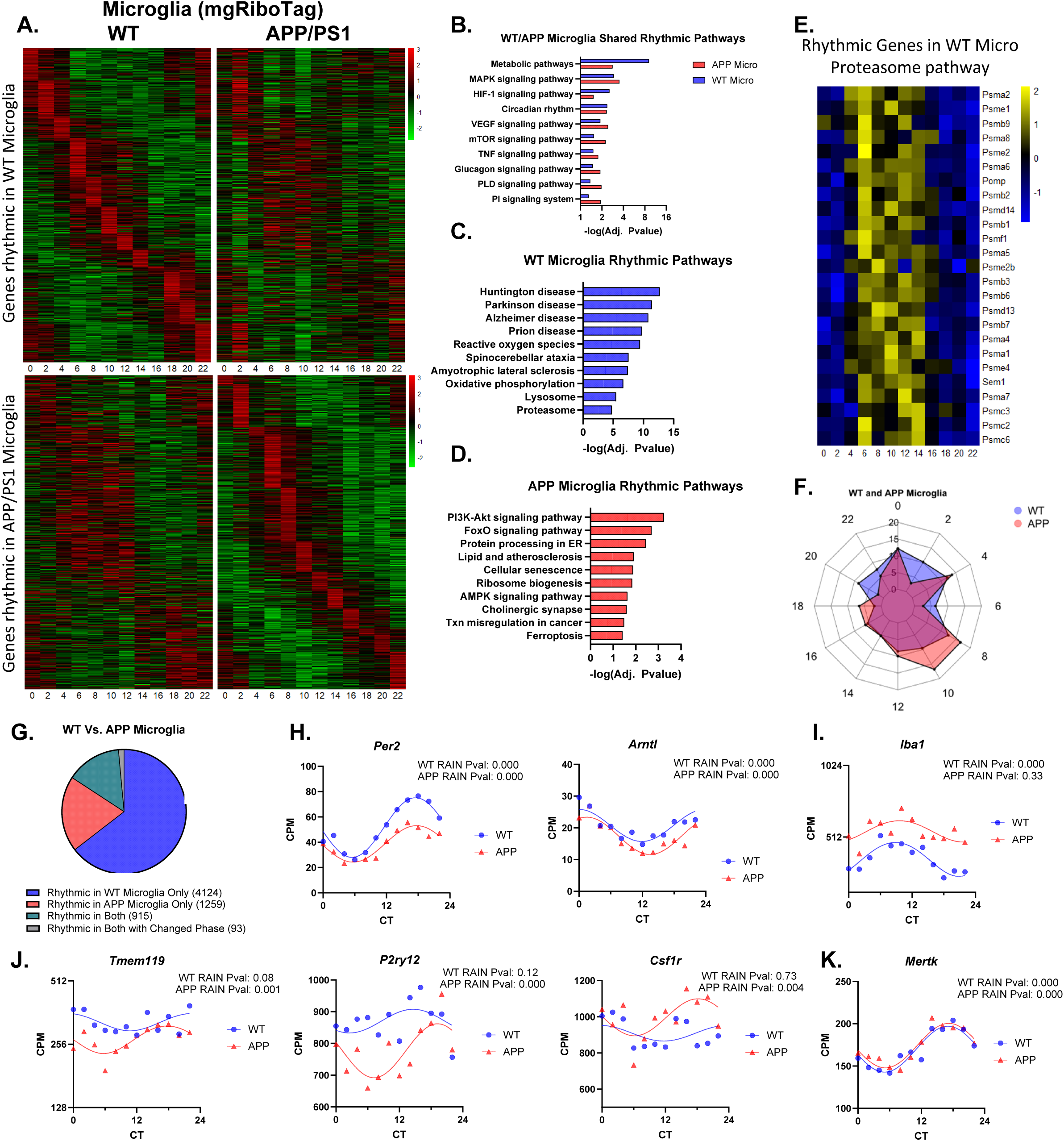
Microglial circadian translatomes and reprogramming in WT and APP/PS1 mice. A. Heatmaps showing transcripts that were rhythmic in WT microglia (upper) and rhythmic in microglia from APP/PS1 mice (lower). In both rows of heatmaps, the genes plotted are in the same order to compare differences in rhythmic expression between mice. B-D. KEGG pathway analysis of microglial transcripts identified as rhythmic (by RAIN analysis) in (B) both WT and APP/PS1 mice, (C) WT mice, or (D) APP/PS1 mice. E. Heatmap showing temporally-coordinated expression of KEGG proteasome pathways genes in microglia from WT mice. F. Radar plot showing acrophase distributions of rhythmic microglial transcripts from WT (pink) or APP/PS1 (blue) mice. G. Pie chart depicting the number of transcripts that gained or lost rhythms in different datasets, based on compareRhythms analysis of rhythmic transcripts in microglia. Total of 6399 were identified as rhythmic by RAIN analysis across all datasets. H-K. Graphs showing circadian expression patterns of transcripts from microglia from WT (blue) or APP/PS1 (red) mice. H. Core clock genes *Per2* and *Arntl* (*Bmal1*) remained rhythmic. I. Iba1-encoding gene *Aif1* lost rhythmicity in APP/PS1. J. Microglial homeostasis markers *Tmem119, P2ry12*, and *Csfr1* gained rhythmicity in APP/PS1. K. Phagocytosis receptor *Mertk* remained rhythmic. Adjusted P values from RAIN are shown. Each datapoint is the average of two mice, each from separate experiments.

Our *compareRhythms* analysis of microglia translatomes showed a larger number of rhythmic transcripts expressed in microglia in general (5039 genes) and found that 82% of these genes (4124) lost rhythmic expression in APP/PS1 mice (Fig. 4G). A smaller set of transcripts (1259 genes) gained rhythmic expression in microglia of APP/PS1 mice (Fig 4G). Similar to astrocytes, we found that circadian clock genes remained rhythmic in both WT and APP/PS1 microglia, suggesting that these differences in rhythmic gene expression were not due to a dysfunctional core clock, though the amplitude of some core clock genes was a mildly blunted in APP/PS1 microglia (Fig 4H and S3F, G). We also found that a number of canonical microglia markers were rhythmic in at least one condition. For example, the microglial gene *Aif1*, which encodes the marker Iba1, was rhythmic in WT microglia, but lost rhythmic expression in APP/PS1 microglia (Fig 4I). Conversely, the homeostatic microglia markers *Tmem119* and *P2ry12*, as well as the disease-associated microglial (DAM) marker *Csf1r,* were non-rhythmic in WT microglia but gained rhythmic expression in APP/PS1 mice (Fig 4J). Many disease-associated microglia (DAM) genes oscillated in microglia in WT mice, but had blunted rhythms in microglia in APP/PS1 mice, including *Tyrobp*, *Ctsb*, *Ctsd*, *Cd9*, *Lpl*, and *Lyz2* (Fig S4A) ^32^. Several lysosomal/proteostatic genes including *Cd68*, *Lamp1,* and *Ctsl* followed a similar pattern (Fig. S4B). We also found that some genes were highly rhythmic in both WT and APP/PS1 microglia, such as the TAM phagocytic receptor *Mertk*, which is critical for microglia engagement of amyloid plaques (Fig. 4K)^33^. Thus, transcripts related to microglial homeostasis and activation state are rhythmic under basal conditions, but these and many other transcripts lose rhythmicity in the presence of amyloid plaques.

We were interested to find that many of the disease-related KEGG pathways (including Alzheimer’s disease) were rhythmic in microglia from WT mice but not APP/PS1 (Fig. 4C, D). To understand if circadian rhythms in gene expression might broadly influence AD-related risk gene expression, we examined the expression of transcripts which had been identified in the most recent AD GWAS as having loci with SNPs (85 total)^34^. We found that transcripts associated with approximately half of all AD GWAS loci were rhythmically expressed exclusively in WT microglia and that many of these genes were non-rhythmic in APP/PS1 microglia (Fig. S5A, B). We also found a substantial number of AD GWAS genes were rhythmic in WT bulk cortical tissue (Fig S5B). Importantly, >80% of these AD-related genes were rhythmic in at least one of our datasets. These data suggest that transcripts related to neurological disease are highly rhythmic in microglia from WT mouse cortex, and that many of these genes seem to lose rhythms in the setting of amyloid plaque pathology.

### Aging induces unique changes to the astrocyte circadian translatome which enhance autophagy and mTOR pathways

We next asked if circadian reprogramming in glia represents a generic response to stress or inflammation, or if reprogramming is truly context-dependent. To test this, we examined aged mice, which are known to develop neuroinflammation, but do not have amyloid plaques. Here, we used the same WT AstroTRAP and mgRiboTag mice, but aged them to 22 months of age (similar to 75 human years)(Fig. 5A). Due to attrition of mice during the aging process, our aged cohorts only had enough mice to sample tissue every four hours over a 24 hour period. Thus, we could not directly compare rhythmic transcripts in young vs. aged mice. Also, rhythmic genes in the aged datasets were defined as having a less stringent RAIN adjusted P value of <0.05 due to the lower number of circadian time points. We first examined rhythmic astrocyte-enriched transcripts in aged mice. Qualitatively, we observed that many transcripts which were rhythmic in young astrocytes (from Fig. 3) were not identified as rhythmic in aged astrocytes, and conversely a number of transcripts were rhythmic in aged astrocytes which were not identified in young ones. (Fig. 5B and S6A). Interestingly, transcripts which were rhythmic in aged astrocytes had very little overlap with the genes that were rhythmic in APP/PS1 astrocytes (Fig 5B, S6A). This suggests that astrocyte genes can gain and lose rhythms in a context-dependent manner. We analyzed KEGG pathways that were rhythmic in aged astrocytes and found a number of pathways that were not identified in WT or APP/PS1 astrocytes (Fig 5C). These included phagocytosis, endocytosis, autophagy, and mTOR signaling. Several rhythmic pathways were similar between young and aged astrocytes, including spliceosome, insulin signaling, and circadian rhythms. FoxO signaling was rhythmic in both APP astrocytes and aged astrocytes, suggesting that this is a conserved response to stress. Plotting rhythmic genes involved in the KEGG endocytosis pathway in aged astrocytes, we were surprised to find that this pathway was largely arrhythmic in astrocytes of younger WT and APP/PS1 mice but showed a temporally-coordinated rhythm in aged astrocytes (Fig. 5D). Examining individual transcripts across all three conditions (young WT, young APP, and aged WT), we found that components of the core circadian clock maintained rhythmicity in aged astrocytes, suggesting that aging did not disrupt the clock in these cells (indeed, absolute expression was actually higher for some transcripts such as *Arntl* and *Per2*, but not *Ciart* Fig. 5E and S6B). In agreement with our pathway analysis, several genes involved in autophagy (*Atg10, Pdpk1, Ulk1*, Fig. 5F) were both increased in expression and were rhythmic only in astrocytes from aged mice. The *Mtor* transcript itself also followed this pattern (Fig. 5G), which is surprising as MTOR activity generally inhibits autophagy. These data suggest a unique response of the circadian translatome to aging (as compared to amyloid). This indicates that aged astrocytes maintain a functional core clock, but that the astrocyte circadian translatome is reprogrammed in a manner which is distinct from that seen in APP/PS1 mice.

**Fig. 5.**
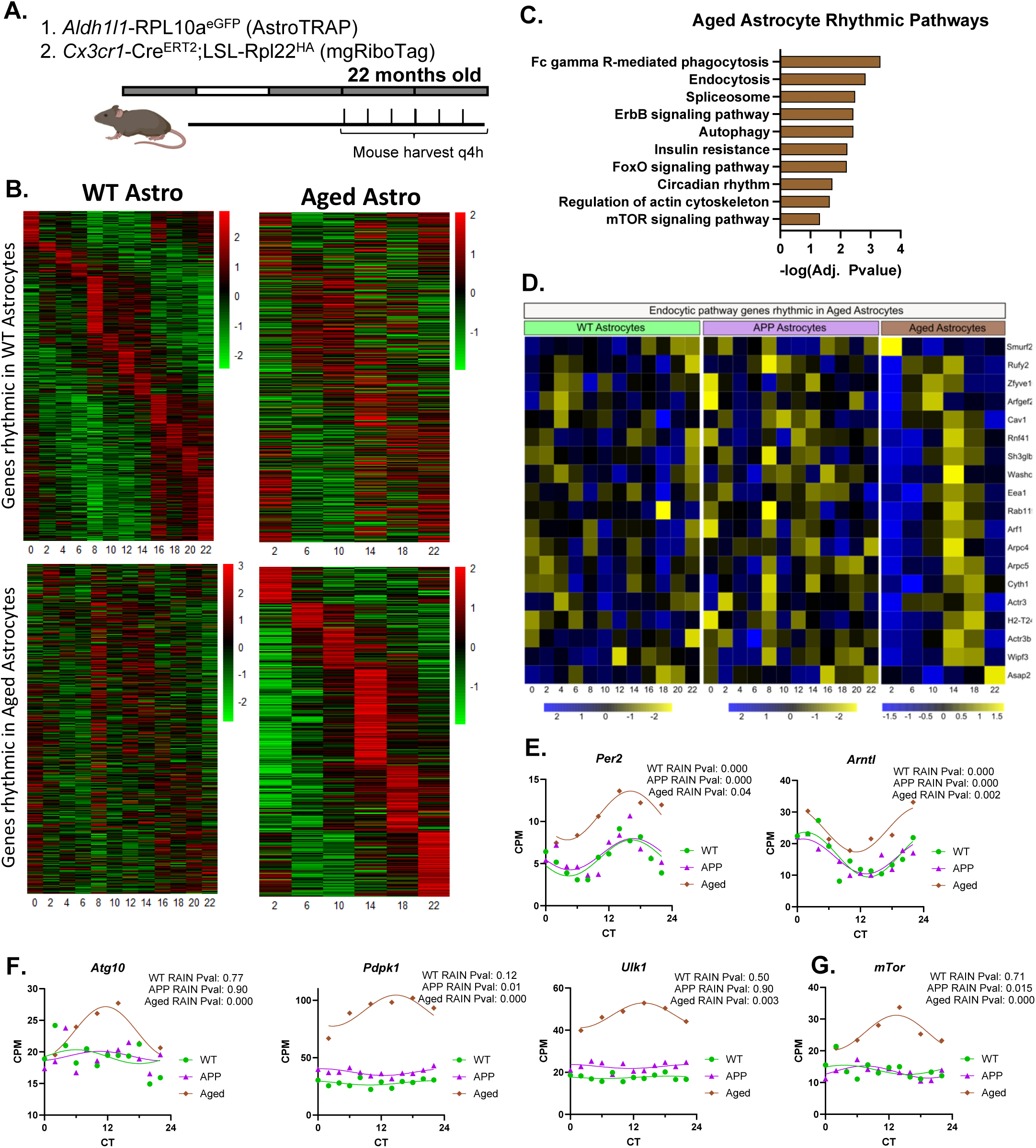
Aging induces rhythmic expression of autophagy and mTOR pathways in astrocytes. A. Schematic of mouse lines and lighting schedule for aging experiment. B. Heatmaps showing transcripts that were rhythmic in astrocytes from young WT mice (upper) and rhythmic in aged mice (lower). In both rows of heatmaps, the genes plotted are in the same order to compare differences in rhythmic expression between mice. C. KEGG pathway analysis of astrocyte transcripts identified as rhythmic (by RAIN analysis) in aged mice. D. Heatmaps showing temporally-coordinated expression of KEGG endocytic pathway genes only in astrocytes from aged mice. E-G. Graphs showing circadian expression patterns of astrocyte transcripts from young WT (green), APP/PS1 (purple), or aged (brown) mice. E. Core clock genes *Per2* and *Arntl* (*Bmal1*) remain rhythmic but are induced in aged astrocytes. F,G. Autophagy genes *Atg10, Pdpk1,* and *Ulk1* (F), as well as *Mtor* (G) are induced and gain rhythmicity in aged astrocytes. Adjusted P values from RAIN are shown. Each datapoint is the average of two mice, each from separate experiments.

### Aging disrupts circadian oscillations in core clock genes and metabolic pathways in microglia

We performed similar circadian analysis on microglia-enriched transcripts from 22mo aged mice and found that aged microglia also dramatically reprogram circadian gene transcription compared to young WT microglia (Fig 6A). Similar to astrocytes, we found that rhythmic pathways in aged microglia were mostly different than those previously seen in WT or APP/PS1 microglia, suggesting microglia genes also gain and lose rhythms in a context-dependent manner (Fig S6C). Pathway analysis of rhythmic transcripts in aged microglia identified several metabolic pathways (TCA cycle, insulin resistance, AMPK signaling), endocytic/proteolytic functions (endocytosis, ubiquitin-mediated proteolysis, peroxisome), and one disease pathway (Parkinson Disease) (Fig. 6B). Interestingly, metabolism, one of the most prominent rhythmic pathways in WT and APP/PS1 microglia, was not identified as rhythmic in aged microglia, suggesting a loss of circadian metabolic control specifically in aged microglia. Heatmap analysis of metabolic pathway transcripts supports this concept (Fig 6C). While the circadian rhythms pathway was identified as rhythmic in aged microglia (Fig. 6B), we were surprised to find that several core circadian clock genes had a dampened expression level and amplitude in aged microglia, including *Arntl*, *Nr1d1*, *Per2* and *Ciart*, the last two of which were not rhythmic in aged microglia (Fig 6D). Contrary to what we found in aged astrocytes, this suggests that core clock function could be dampened in aged microglia. The endocytosis gene *Rab5c, mitochondrial genes Ndufa10,* and the proteasome subunit gene *Psmd11* were increased in expression and rhythmic only aged microglia, again suggesting circadian regulation of a unique proteostatic response in aging (Fig 6E and S6D). Some transcripts maintained rhythmicity in WT, APP/PS1 and aged microglia, but had dampened expression only in aged microglia, including the low-density lipoprotein receptor gene *Ldlr* while other genes completely lost rhythms in aged microglia, like transcription factor *Mafg* (Fig 6F). These data suggest that microglia lose rhythmic expression of many of their genes during normal aging, including a dampening of core circadian clock genes and metabolic pathways.

**Fig. 6.**
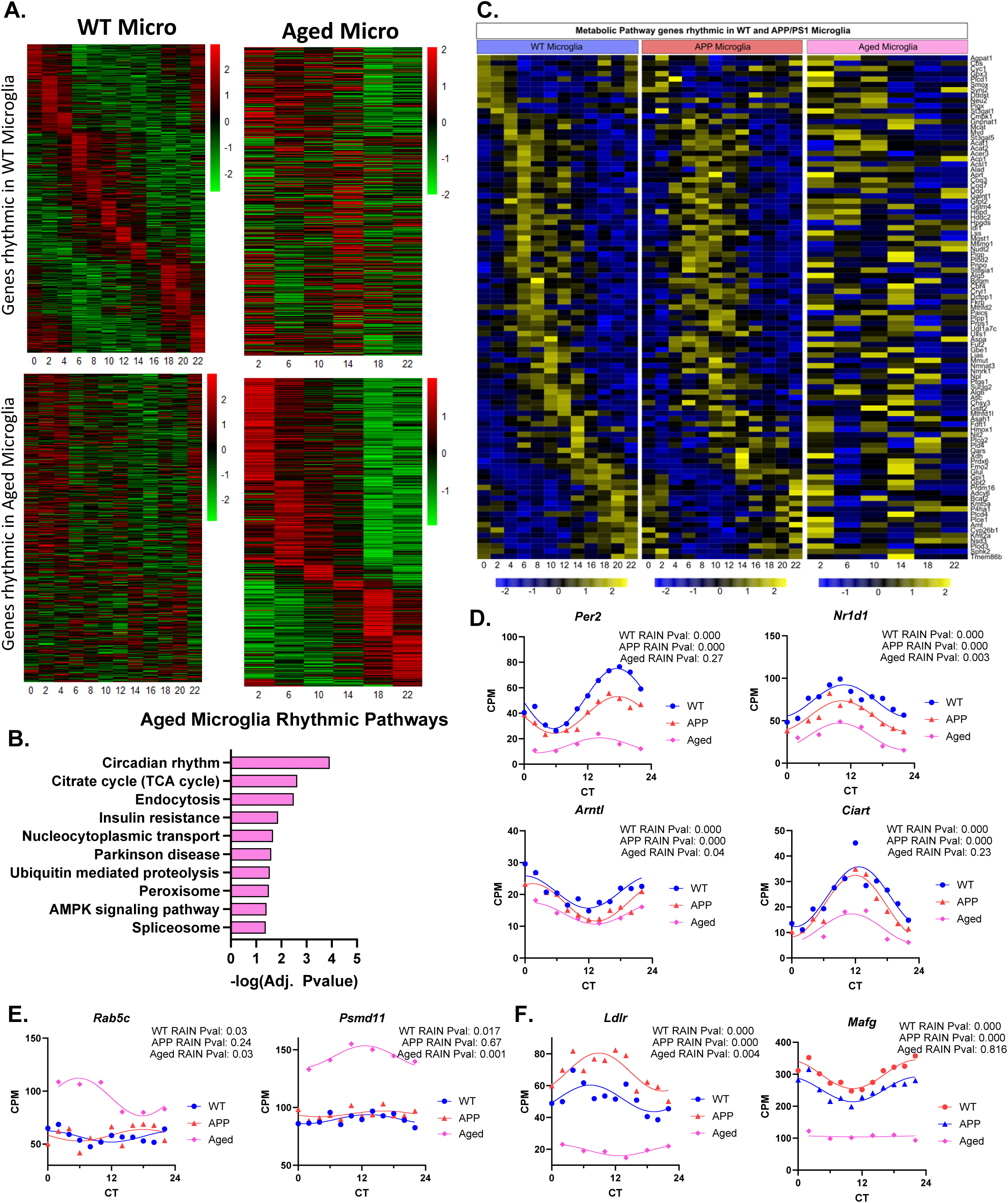
Aging suppresses microglial core clock oscillation and circadian gene expression. A. Heatmaps showing transcripts that were rhythmic in microglia from young WT mice (upper) and rhythmic in aged mice (lower). In both rows of heatmaps, the genes plotted are in the same order to compare differences in rhythmic expression between mice. B. KEGG pathway analysis of microglial transcripts identified as rhythmic (by RAIN analysis) in aged mice. C. Heatmaps showing temporally-coordinated expression of KEGG metabolic pathway genes in microglia from young WT, APP/PS1, and aged mice. D-F. Graphs showing circadian expression patterns of transcripts from microglia from WT (blue) or APP/PS1 (red) mice. D. Core clock genes *Per2*, *Arntl* (*Bmal1*), *Nr1d1,* and *Ciart* remained rhythmic but were blunted and suppressed in aged microglia. E. Endosomal trafficing gene *Rab5c* and proteasome subunit *Psmd11* are induced and gain rhythmicity in aged microglia. F. The lipoprotein receptor *Ldlr* and transcription factor *Mafg* lose rhythmicity in aged microglia. Adjusted P values from RAIN are shown. Each datapoint is the average of two mice, each from separate experiments.

### Disease associated signatures in glia depend on the time of day

Numerous groups have attempted to determine gene expression signatures of disease associate microglia and astrocytes in AD models^32, 35, 36^. However, given the dramatic changes in rhythmicity we observe above, it is unclear if such signatures are stable over circadian time. Considering the broad and complex circadian regulation of transcripts in the brain across cell types and disease states, we next sought to understand the potential impact that time of day of mouse harvest has on identifying differential gene expression in microglia and astrocytes between WT and APP/PS1 mice. We first enriched for cell-specific transcripts by including only transcripts that were significantly upregulated in our datasets when compared against the Pre-IP datasets. We then binned time points together to represent general times-of-day that a morning person or an evening person might sacrifice mice in the lab (6-10am are AM, vs. 4pm-8pm are PM) and compared differential gene expression between WT and APP/PS1 microglia samples. Differential gene expression analysis of mice sacrificed in the AM (6-10am timepoints) identified 506 upregulated genes in microglia from APP/PS1 mice (adjusted pvalue <0.05 and fold change >1.5) (Fig 7A), while identical analysis of mice sacrificed in the PM (4-8pm), revealed a ∼20% increase in the number of upregulated genes (627 genes total) (Fig 7B). Restricting analysis to differentially-expressed genes (adjusted pvalue <0.05 and fold change >/< 1.5), and comparing fold changes (WT vs. APP/PS1) in the AM versus PM groups, we found that genes significantly change their expression patterns based on the time of day of harvest, influencing differential gene expression analysis. This is especially prominent if we focus on genes with lower fold changes (Fig 7C). Examining individual transcripts, we observed several important microglial genes that are only significantly differentially expressed at one time of day. For example, microglia transcription factor, *Spi1,* interferon-inducible protein *Aim2*, and phagocytic regulator *Cd209a* were only differentially expressed if tissue was harvested at a specific time of day (Fig. 7D and E). A number of other genes, including *Ldlr, Spi1*, and *Col22a1* were also only differentially expressed at one time of day (Fig S7A and S7B). When analyzing the significantly upregulated genes in the AM vs. PM time bin, we found that more than 25% of the identified genes could be specific for a particular time of day (S7E).

**Fig. 7:**
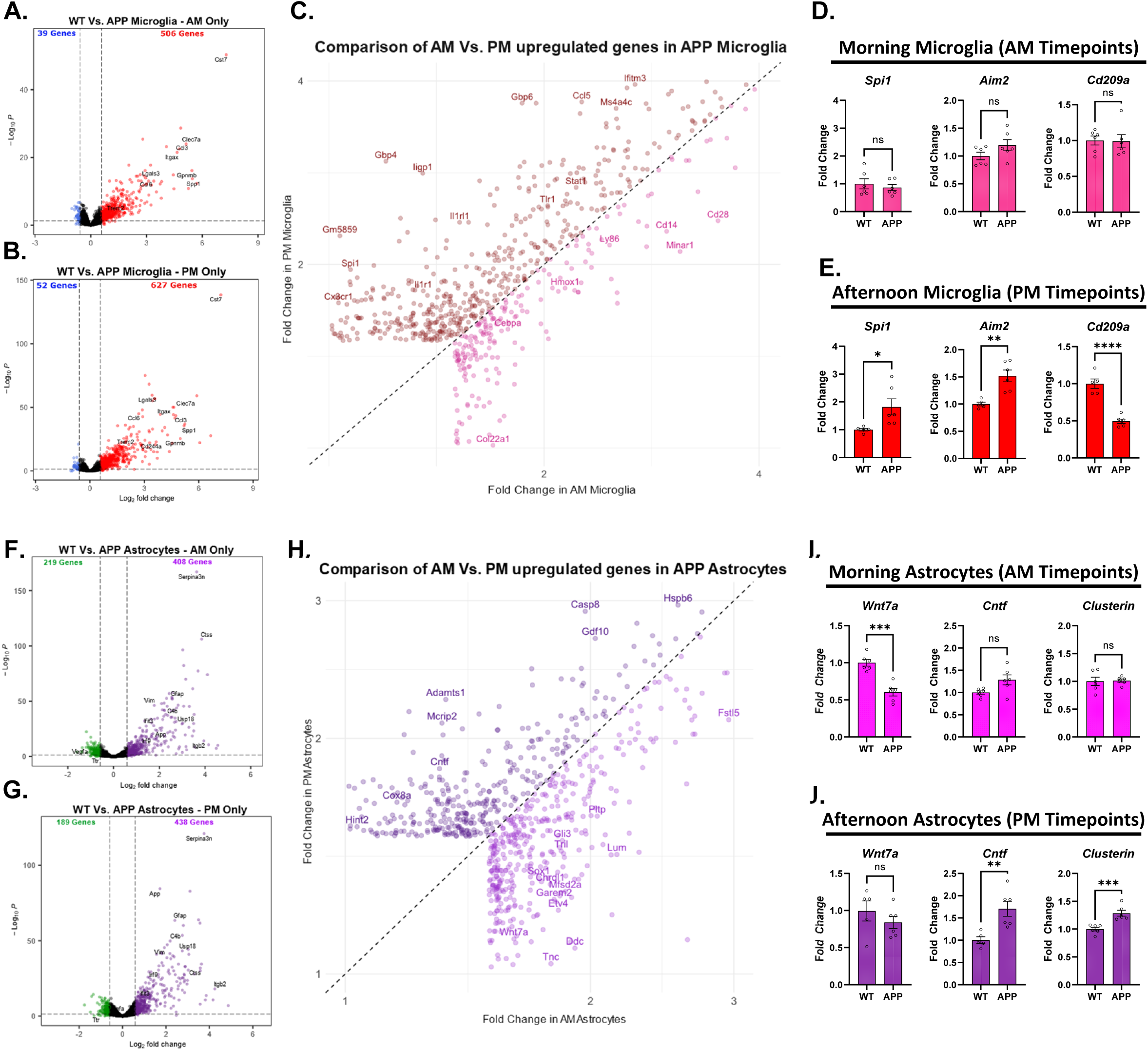
Time-of-day of tissue harvest influences differential gene expression in APP/PS1 mice. A & B. Volcano plots showing differential gene expression in microglia between WT and APP/PS1 mice harvested during the morning (AM) hours (A) or evening (PM) hours (B). 506 DEGs were identified in AM, 627 in PM. C. Comparison of fold change of induction of DEGs in microglia in WT vs. APP/PS1 mice harvested in AM or PM. Brown circles indicate more expression in PM; pink circles indicate more expression in AM. D & E. Genes plotted from microglia in the AM time bin (D) or PM time bin (E) including myeloid transcription factor *Spi1*, interferon-inducible protein *Aim2*, and *Cd209a*, known to regulate phagocytic activity. Volcano plots showing differential gene expression in astrocytes between WT and APP/PS1 mice harvested during the morning (AM) hours (F) or evening (PM) hours (G). 438 DEGs were identified in both AM and PM, though not the same 438. H. Comparison of fold change of induction of DEGs in astrocytes in WT vs. APP/PS1 mice harvested in AM or PM. Dark purple circles indicate more expression in PM; light purple circles indicate more expression in AM. I & J. Genes plotted from astrocytes in the AM time bin (I) or PM time bin (J) including, WNT pathway gene *Wnt7a*, ciliary neurotrophic factor *Cntf*, and AD GWAS gene, *Clu* (Clusterin). In C and H, only genes with fold change <3 were examined-all genes with large fold changes (>3 fold) were identified as DEGs in both AM and PM datasets. Graphs in (D,E,I and J) show mean ± SEM. **p < 0.01, ***p < 0.001 by unpaired two-tailed t test

We performed a similar analysis on WT and APP astrocyte transcripts, binning the same morning and evening time points. Similar to microglia, there were significantly more differentially expressed genes in astrocytes in the PM group (438 genes) than AM (406 genes, Fig 7F, G). Comparing the fold changes from significantly dysregulated genes in astrocytes from APP/PS1 mice sacrificed in the AM vs. PM, we found many genes that had different fold changes between morning and afternoon (Fig 7H). This effect was more prominent in genes with smaller fold change. Examining individual transcripts, we observed several astrocyte genes that are only significantly differentially expressed at one time of day. For example, WNT pathway gene *Wnt7a*, *Cntf* (encoding ciliary neurotrophic factor), and AD GWAS gene *Clu* (encoding Clusterin/ApoJ) were only differentially expressed at a specific time of day (Fig. 7J). A number of other transcripts were also differentially expressed in either the AM or PM time bin, including *Pparα*, *Ldha*, and AD GWAS gene *Adamts1* (Fig S7C and S7D). Similar to microglia, we found that more than 25% of significantly upregulated genes identified were specific to the time of day the mouse was sacrificed. Thus, time-of-day of mouse harvest influences the differential gene expression that is observed in astrocytes and microglia.

Finally, we generated a website which provides a publicly-available interface to search and visual our dataset. The website provides automatic data visualization of gene expression across timepoints for each dataset, average absolute expression level of the gene of interest in each dataset, and P values for each dataset generated by RAIN. This resource will allow researchers to query the circadian expression patterns of any gene across timepoints, cell types, and conditions, and is available at https://musieklab.shinyapps.io/Glial_Circadian_Translatome/. Typical output from the site is shown in Fig. S8.

## Discussion

Here we have used TRAP/RiboTag-RNAseq to elucidate the circadian translatomes of astrocytes and microglia in healthy mouse brain, and in response to amyloid plaque pathology or aging. Our data reveal that astrocytes and microglial have robust and unique circadian translatomes, that these translatomes change dramatically in the setting of amyloid pathology or aging, and that changes are cell type-specific, and context-dependent. The core circadian clock was generally robust in the setting of amyloid plaque pathology in bulk cortex, astrocytes, and microglia, though downstream rhythms in gene expression underwent robust circadian reprogramming. However, aging caused blunting of core clock gene rhythms in microglia, but not astrocytes. Many AD-related genes showed circadian rhythms in one cell or condition, with several gaining rhythmicity in response to amyloid plaques. Finally, the landscape of differentially expressed genes in APP/PS1 mice with amyloid plaques varies considerably based on the time of sacrifice of the mice, with more DEGs being identified in the evening. Our results not only create a valuable resource for other investigators to study circadian gene expression in glial cells in the brain in health and disease, but also illustrates the broad and complex effects of the circadian system on critical pathways related to microglia and astrocyte function, particularly in the context of AD pathology.

Our findings here build on numerous studies which show that genetic disruption of the circadian clock in glial cells has marked effects on brain homeostasis and AD-related pathology. Cell type specific deletion of the key circadian transcription factor *Bmal1* in astrocytes causes astrocyte activation, inflammatory gene expression, and can mitigate tau and alpha-synuclein pathology in mouse models^21, 22, 24^. Microglial *Bmal1* deletion can impair synaptic pruning and accelerate brain aging, but can also suppress IL-6 expression and reduce lesion volume after stroke^20, 38^. Deletion of the core clock gene *Nr1d1* (REV-ERBα) in microglia can promote inflammation, leading to reduced amyloid plaque pathology in 5xFAD mice, but exacerbated tauopathy in P301S PS19 mice^23, 39^. Our dataset presented here should help shed light on the mechanisms by which glial clocks influence brain health. As a specific example, we have previously shown that astrocytic *Bmal1* deletion can induce endolysosomal and autophagic gene and protein expression and enhance astrocytic protein degradation capacity^22, 24^. In our studies here, we identified pathways related to endolysosomal function and autophagy as having circadian rhythmicity in bulk cortex, astrocytes, and microglia in several contexts. In bulk cortex, pathways related to lysosome, autophagy, and mTOR signaling were identified as rhythmic in WT mice, but all lost rhythmicity in APP/PS1 mice. In microglia, both lysosome and proteasome pathways were rhythmic in WT mice but also lost rhythms in APP/PS1 mice. Proteostatic pathways were not highly rhythmic in astrocytes in either young WT or APP/PS1 mice, but autophagy and mTOR signaling were upregulated and gained rhythmicity in aged astrocytes, suggesting that *Bmal1* deletion bears more similarity to astrocyte aging than to amyloid exposure. Similarly, pathways related to endocytosis and ubiquitin-mediated proteolysis gained rhythmicity in aged microglia, suggesting that the circadian clock reprograms to regulate proteostatic functions in glia as a conserved aspect of brain aging. These findings suggest that amyloid pathology may impair circadian regulation of proteostatic functions in the brain, while aging may induce them. This could be a mechanism by which amyloid pathology acts as a catalyst for subsequent tau and alpha-synuclein aggregation in AD. It remains unclear if this increase in circadian rhythmicity of proteostatic genes occurs in response to accumulating proteostatic stress in aged glia, or if this response is beneficial or harmful. It will be critical to understand how circadian reprogramming of proteostatic gene expression occurs in glial, if it plays a key functional role on brain protein aggregation during aging, and if it can be harnessed therapeutically to prevent neurodegenerative pathologies. Furthermore, as therapies are developed for neurodegenerative diseases, it may be important to consider the time of day of administration, as many therapies, such as chemotherapy of brain tumors, has been shown to be more effective when administered at a specific time of day ^40^.

One key implication of our data is that the identification of differentially-expressed genes in a disease state may vary substantially depending on the time of day of tissue harvest. As there are now hundreds of AD-related transcriptomic datasets being used as resources by researchers around the world, time-of-day of tissue harvest should be considered when assessing the differential expression of a given gene, or when trying to combine datasets that may have been harvested at different circadian phases. In many cases the tissue may have been harvested at a variety of timepoints and averaged together, which may create significant noise at obscure changes in gene expression.

In summary, we present a resource for exploring the circadian translatomes of astrocyte, microglia, and bulk cortex in vivo under basal conditions and in the setting of amyloid plaques and aging. Our findings illustrate that circadian rhythms in gene expression are highly dependent on cell type and are reprogrammed in a context-dependent manner, in some cases despite robust core clock oscillation. We find that many transcripts related to metabolism, proteostasis, and Alzheimer’s Disease exhibit rhythmic expression that can be altered by pathology, emphasizing the need to further understand circadian changes in gene expression and cellular function in aging and neurodegenerative conditions.

## Methods

### Sample collection

#### Mice

All animal experiments were approved by the Washington University IACUC and were conducted in accordance with AALAC guidelines and under the supervision of the Washington University Department of Comparative Medicine. Two mice were used for every time point, one male and one female to avoid any sex bias in gene expression or sex-specific circadian rhythms. Some samples were excluded from analysis due to poor RNA quality, these timepoints were then analyzed in singlicate: Bulk WT 6pm, WT Microglia 6pm, WT Microglia 10pm, Aged Astrocytes 8pm, and Aged Microglia 8pm. For all samples except the aged samples, mice were 6-7 months old. For all aged samples, mice were 21-23 months old. To pull down astrocyte-specific RNA, Aldh1l1:EGFP/RPL10A mice were used. For microglia-specific RNA, Cx3cr1-Cre^ERT2^-IRES-eYFP^+/−^; Rpl22^HA/+^ were used.

#### Circadian processing of samples

Prior to sacrificing, mice were place in a dark room for 48-72 hours in order to remove any effects of light-induced transcription. One WT and one APP mouse were sacrificed every two hours over a single 24 hour period from either the Aldh1l1:EGFP/RPL10A or Cx3cr1-Cre^ERT2^-IRES-eYFP+/−; Rpl22^HA/+^ genotypes. This was performed a second time to add replicates to each time point. At each time point, mice were injected with fatal plus in the dark. Once the mice were unresponsive, they were perfused with 30mLs of PBS supplemented with 100ug/mL cycloheximide in order to stop ribosomes in place. The brains were then removed, cortices were isolated and frozen at −80°C until processing.

### Ribosome immunoprecipitation and RNA isolation

For processing of ribotag samples, microglia-specific RNA was isolated by first sonicating tissue with 3mLs of homogenization buffer. Samples were centrifuged at 10,000rpm at 4°C for 10 minutes. Supernatant was collected and 10ul of 1mg/mL anti-HA antibody was added (Biolegend Clone 16B12). Samples were placed at 4C on a rotator for four hours. Washed Protein G beads were then added to each tube and incubated overnight. Samples were washed three times with a high salt buffer at 4C using a magnet to collect beads for each wash. Lysis buffer was then added to each sample before isolating RNA using the RNeasy Micro Kit.

For processing TRAP samples, streptavidin Dynabeads were washed and mixed with biotinylated protein L. After 35 minutes, beads were washed and 50ug of anti-eGFP antibodies were added (19C8+19F7 antibodies) for one hour. These bead antibodies were washed 3x before they were used. Frozen brain samples were dounce homogenized in homogenization buffer. The lysed samples were centrifuged at 20,000xg for 10 minutes and supernatants were transferred to new chilled tubes, with bead/antibody complexes. From this tube, 1/10 of the sample was taken and added to 700ul of Trizol for all Bulk samples (Pre-immunoprecipitation of astrocyte-specific RNA). The homogenate and beads were then incubated for 4 hours at 4C. Beads were collected by magnet and washed with salt buffer. After the 5^th^ wash, beads were resuspended in trizol and RNA isolated using RNA Clean and Concentrator kit (Zymo Research).

### RNA Sequencing

Every sample was prepared as follows: Total RNA integrity was determined using Agilent Bioanalyzer or 4200 Tapestation. Library preparation was performed with 10ng of total RNA with a Bioanalyzer RIN score greater than 8.0. ds-cDNA was prepared using the SMARTer Ultra Low RNA kit for Illumina Sequencing (Takara-Clontech) per manufacturer’s protocol. cDNA was fragmented using a Covaris E220 sonicator using peak incident power 18, duty factor 20%, cycles per burst 50 for 120 seconds. cDNA was blunt ended, had an A base added to the 3’ ends, and then had Illumina sequencing adapters ligated to the ends. Ligated fragments were then amplified for 12-15 cycles using primers incorporating unique dual index tags. Fragments were sequenced on an Illumina NovaSeq-6000 using paired end reads extending 150 bases.RNA-seq reads were then aligned and quantitated to the Ensembl release 101 primary assembly with an Illumina DRAGEN Bio-IT on-premises server running version 3.9.3-8 software.

### RAIN analysis

The filterByExpr() function from the EdgeR package was used to filter low counts from each sample group. The *ComBat_seq()* function was used for batch correction (from the sva R package version 3.46.0). CPM’s were calculated using the cpm() function from the EdgeR package. Batch corrected CPM’s were fed into R package rain (version 1.32) to determine circadian gene expression in each dataset. For the majority of samples, genes were considered rhythmic if the FDR adjusted P value from RAIN was <0.01. For all aged samples, genes were considered rhythmic if the FDR adjusted P value was <0.05.

### KEGG pathway analysis

Lists of rhythmic genes from each dataset were identified as described above, then entered into the Database for Annotation, Visualization and Integrated Discovery (DAVID, https://david.ncifcrf.gov/) ^48^. KEGG Pathway analysis was performed and files from this were downloaded. Full list of significant KEGG pathways are provided in the supplement.

### Differential Expression analysis

To identify astrocyte and microglia-specific genes in each dataset, differential expression with DESeq2 (version 1.38.3) was used with each dataset comparing against Bulk sequenced datasets to determine only cell-type specific significantly enriched genes (log fold change > 0 and an adjusted pvalue < 0.05). These cell-specific genes were then used and specific time points were isolated (AM and PM). Differential expression using DESeq2 was then run on AM and PM groups comparing WT Vs. APP samples.

## Data Availability

All raw RNAseq files are available at GEO, accession number GSE261698. Data may also be explored and downloaded at https://musieklab.shinyapps.io/Glial_Circadian_Translatome/.

## Author Contributions

The experimental plan was developed by PWS, JDD, JDF, and ESM. The experiments were carried out by PWS, DS, SK, and JL. Data analysis was performed by PWS, SF, HH, RCA, and ESM. The website was developed by PWS and SF. PWS, SF, and ESM prepared the manuscript. Funding was provided by ESM, JDF, and JDD.

## Acknowledgements

This work was funded by NIA grant R01AG054517 (ESM and JDF) and NINDS grant R01NS102272 (JDD). PWS was funded by NIA grant T32AG058518. DS was supported by NIH grant R00AG061231 and the Rockford Draper Early-Career Faculty Development Award. We thank the Washington University Genome Technology Access Center at the McDonnell Genome Institute (GTAC@MGI) for carrying out RNAseq.

---

**Fig. S1.**
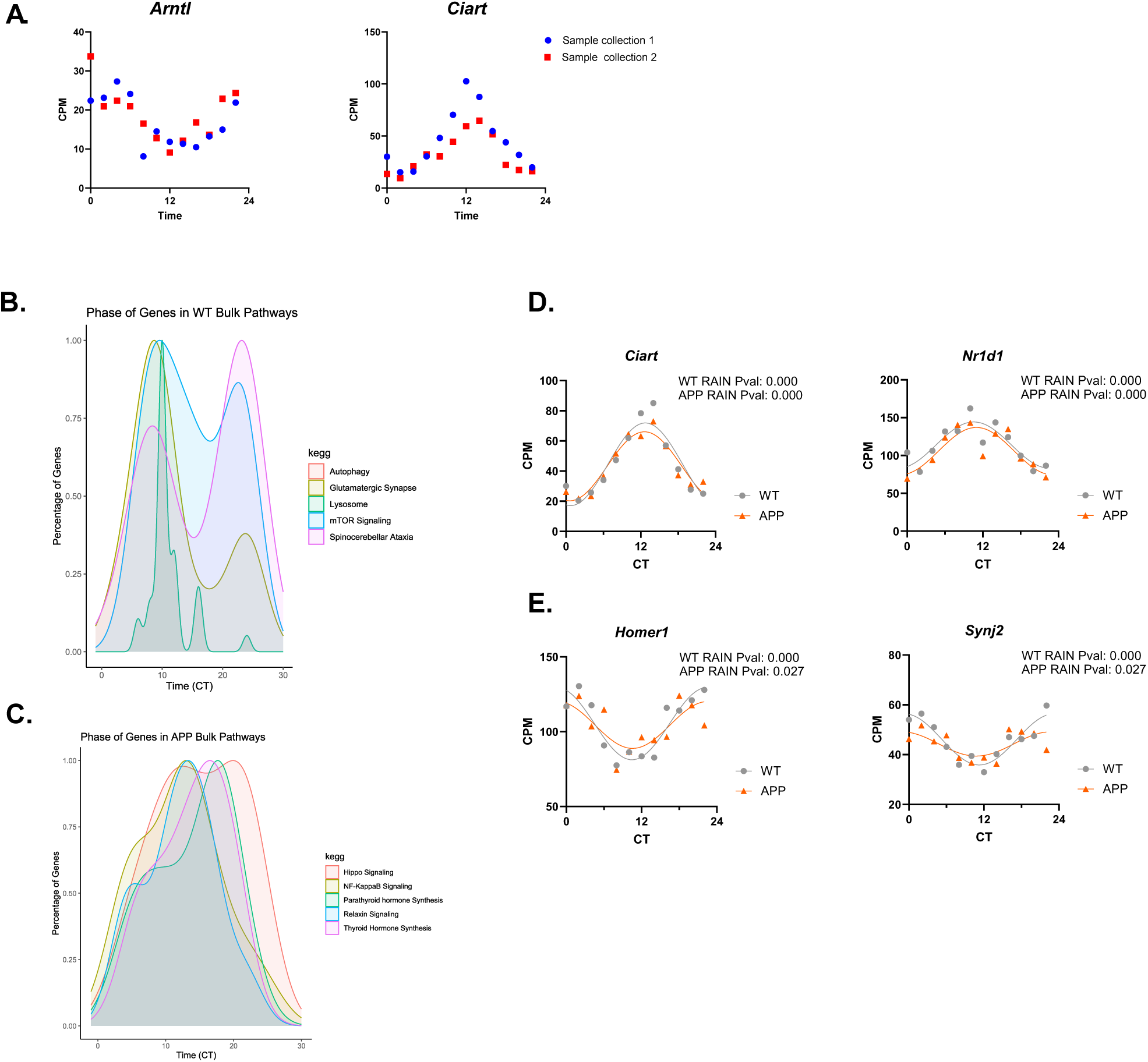
: Additional analysis of circadian gene expression in bulk cortex tissue. A. Graphs depicting the reproducibility of circadian expression of core clock genes *Arntl* (Bmal1) and *Ciart* between experimental replicates. Blue circles are data from mice harvested in the first experiment, and red are from the second circadian harvest. Each datapoint represent one mice. These duplicates were averaged after batch correction at each timepoint to generate the data in all other figures. B & C. Graphs showing the distribution of acrophase values of rhythmic transcripts in bulk cortex from WT (B) and APP/PS1 (C) cortex. Transcripts are grouped by KEGG pathway. D. Circadian expression of core clock genes Ciart and Nr1d1 in bulk cortex from WT (grey) and APP/PS1 (orange) mice. E. Circadian expression of synaptic genes Homer1 and Synj2 in bulk cortex from WT (grey) and APP/PS1 (orange) mice. For D and E, RAIN P values are shown, and each datapoint represents the average of two mice harvested in separate experiments.

**Fig. S2.**
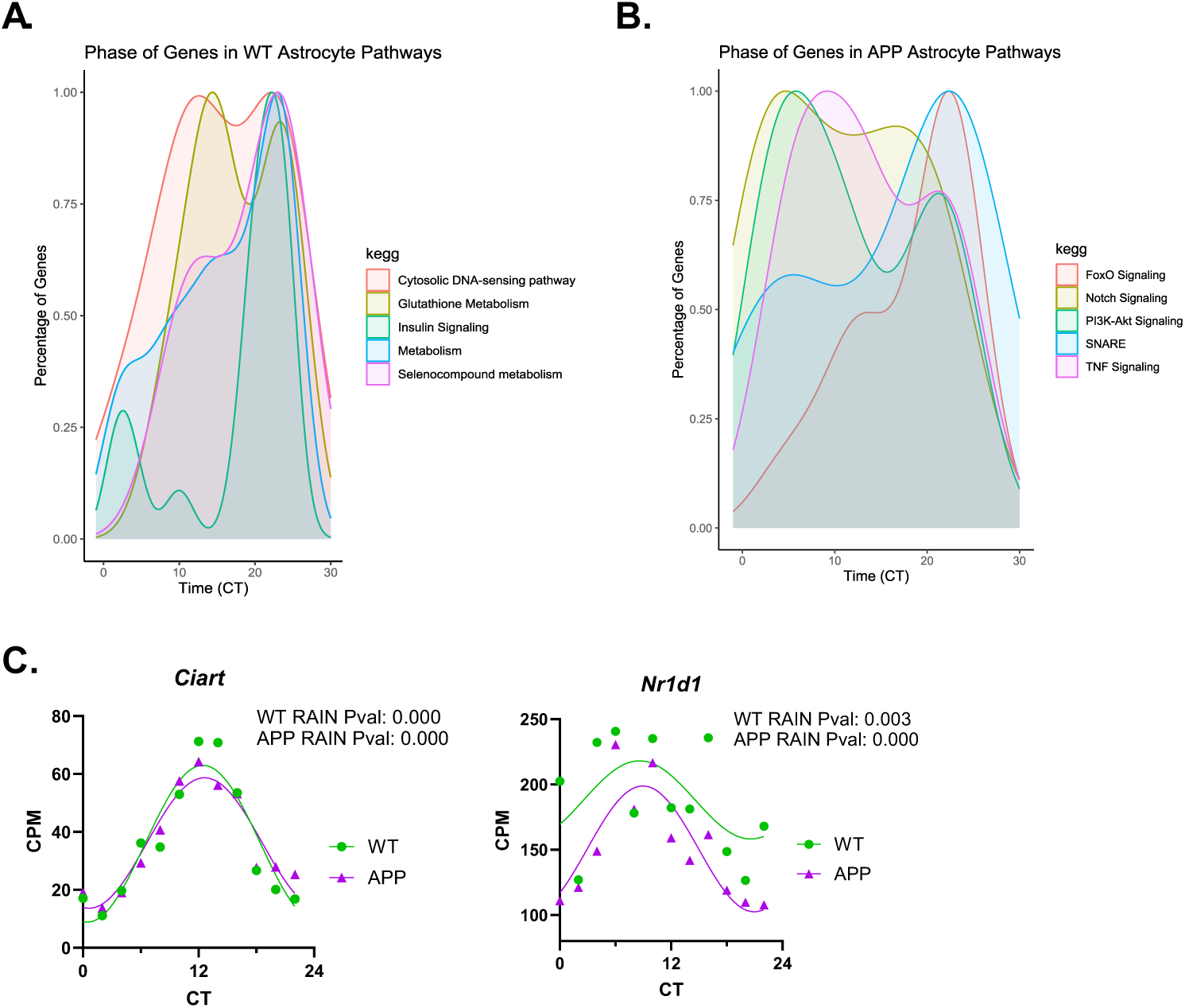
: Additional analysis of circadian gene expression in astrocyte. A & B. Graphs showing the distribution of acrophase values of rhythmic transcripts in astrocytes from WT (A) and APP/PS1 (B) cortex. Transcripts are grouped by KEGG pathway. C. Circadian expression of core clock genes *Ciart* and *Nr1d1* in bulk cortex from WT (green) and APP/PS1 (purple) mice. RAIN P values are shown, and each datapoint represents the average of two mice harvested in separate experiments.

**Fig. S3:**
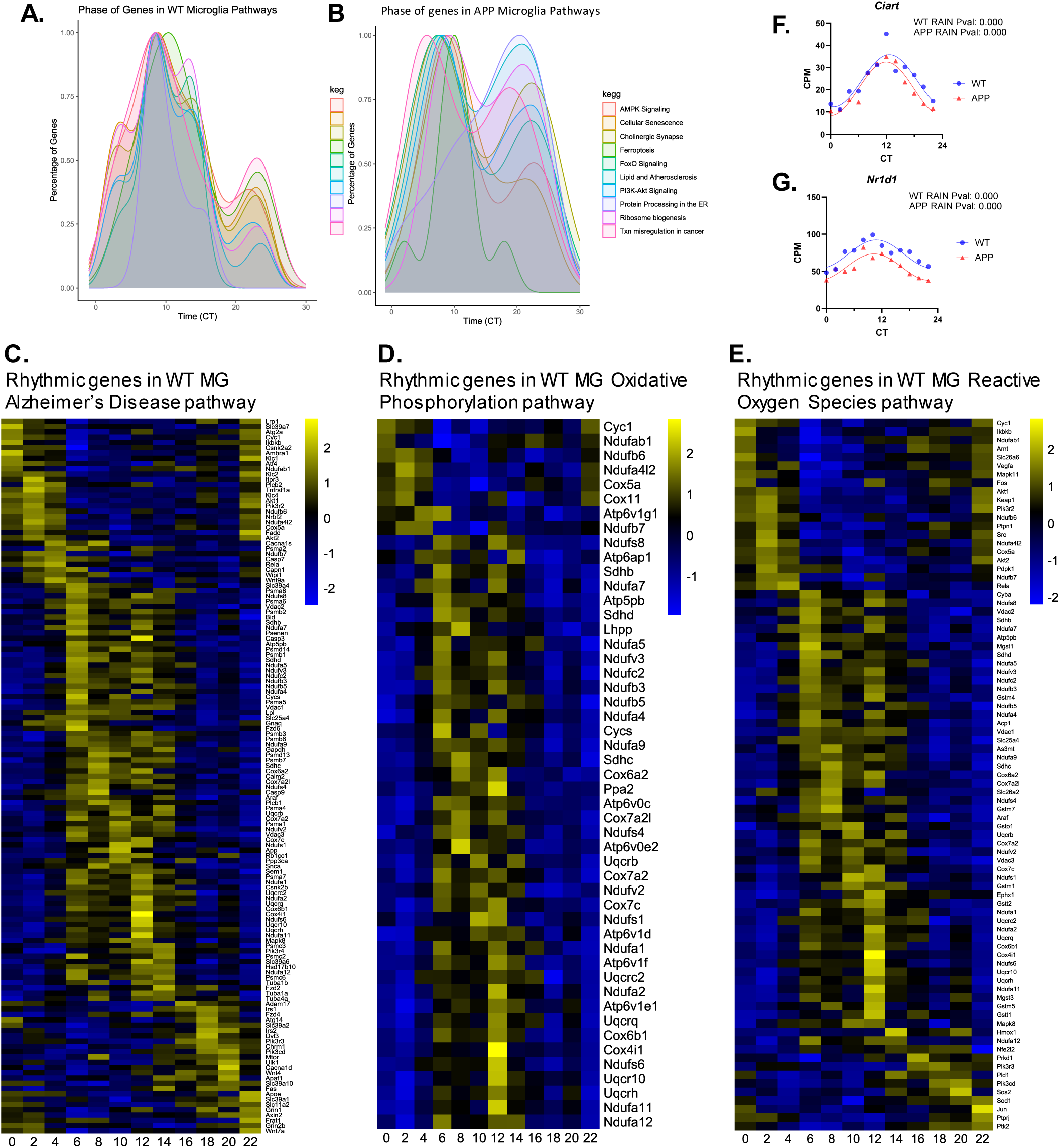
Additional characterization of circadian rhythms in gene expression in microglia. A & B. Graph showing the acrophase of rhythmic genes in microglia from WT (A) and APP/PS1 (B) mice. Genes are separated by KEGG pathway. C-E. Heatmaps showing temporally-coordinated gene expression in microglia (MG) from WT mice for the Alzheimer’s Disease (C), oxidative phosphorylation (D), and reactive oxygen species (E) KEGG pathways.. F & G. Rhythmic expression of clock genes *Ciart* (F) and *Nr1d1* (G) showing preserved rhythmicity in APP/PS1 mice (APP, red). P values from RAIN analysis are shown.

**Fig. S4:**
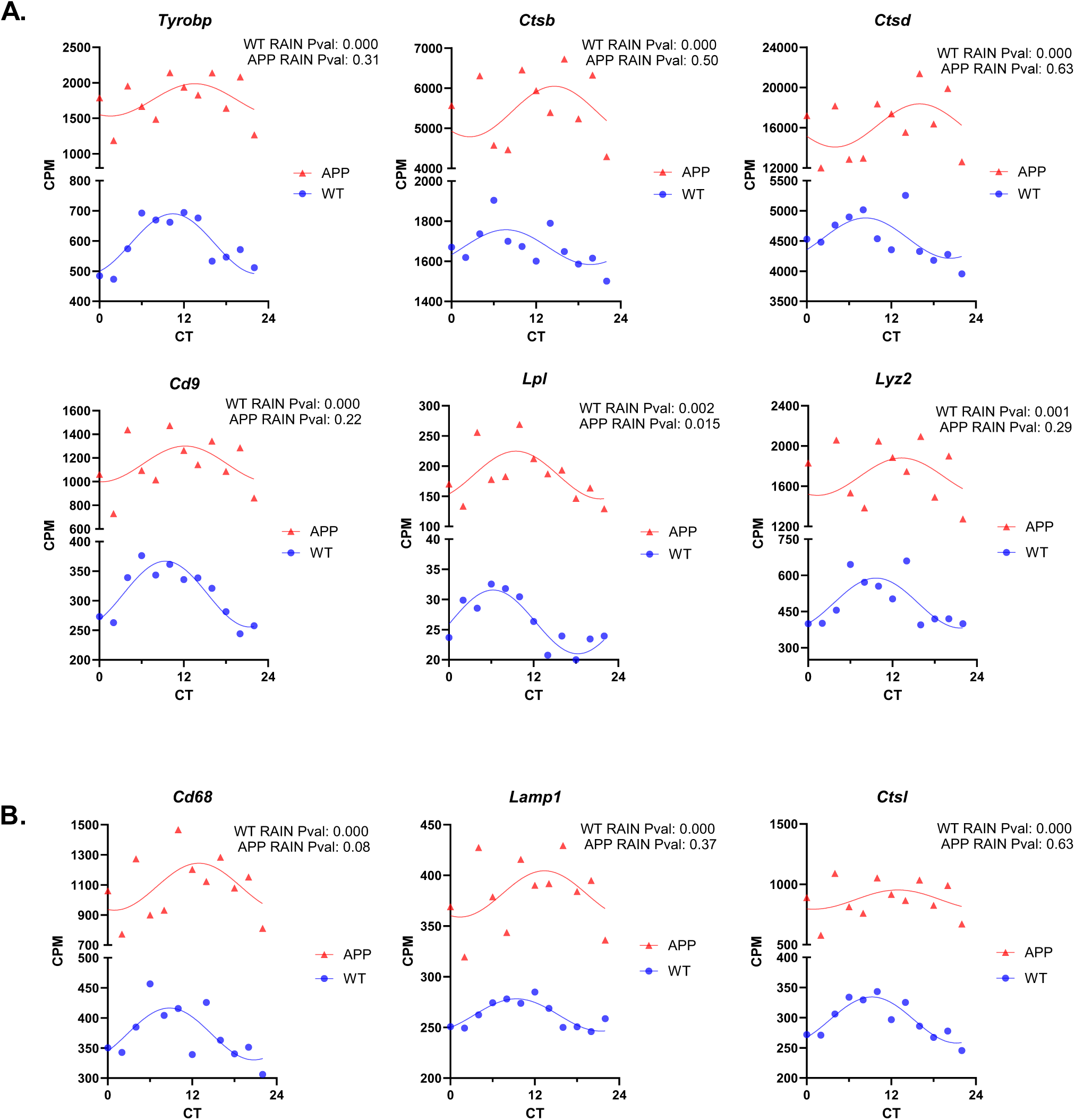
Circadian rhythms in DAM and phagolysosomal genes in microglia in healthy mouse brain. A. Graphs depicting circadian rhythms in expression of disease-associated microglial (DAM) activation genes. Adjusted P values from RAIN are shown. B. Graphs depicting circadian rhythms in the microglial phagosome marker gene Cd68, the lysosomal gene Lamp1, and the protease cathepsin L (*Cstl*) in WT mouse brain. Adjusted P values from RAIN are shown. Of all transcripts shown in A and B, only *Lpl* (P=0.015) was rhythmic in APP/PS1 mice. Note that most graphs have a 2-segment Y axis due to upregulation of transcripts in APP/PS1 mice.

**Fig. S5:**
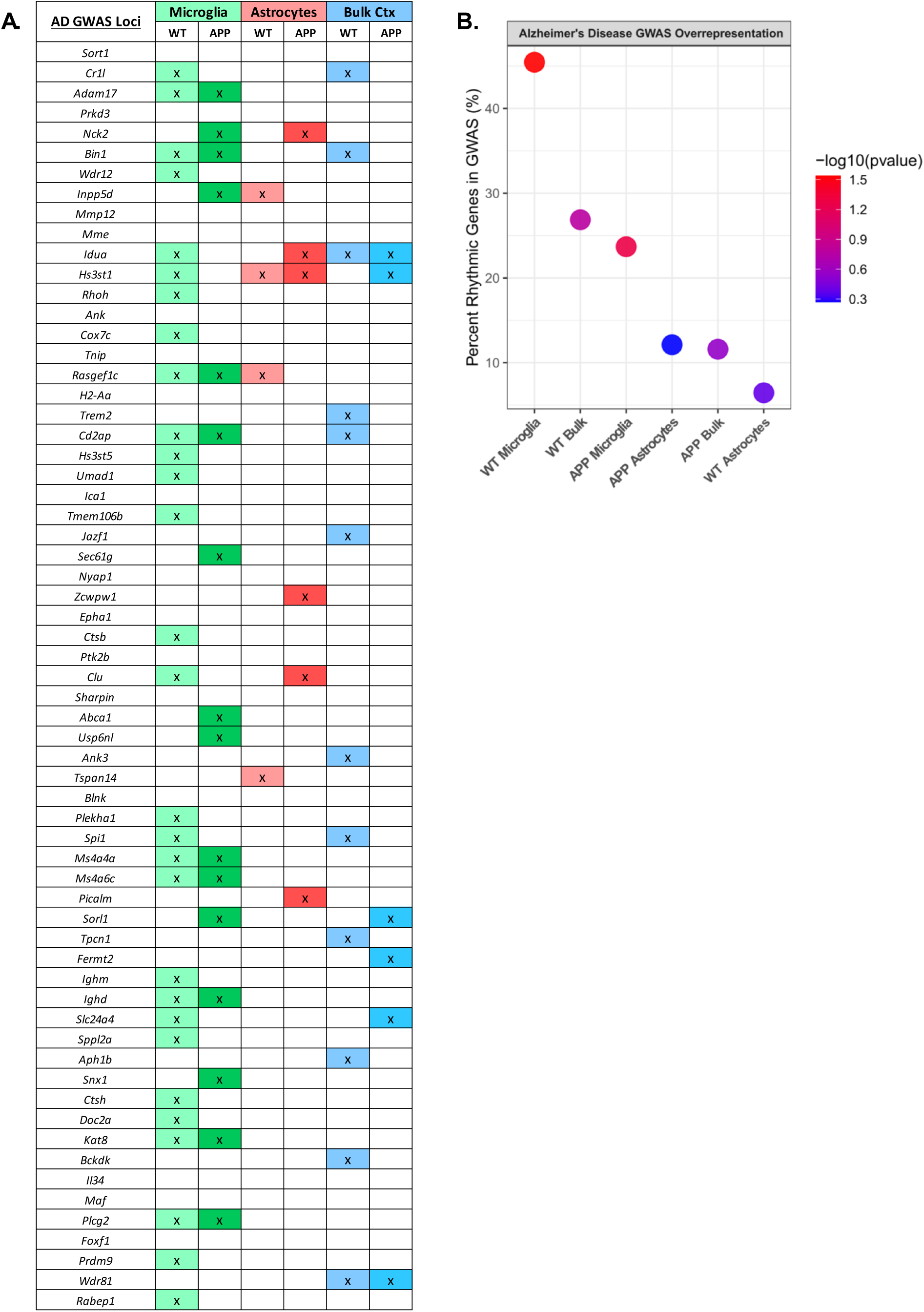
Circadian rhythms in AD GWAS genes. A. Table listing current AD risk genes as identified by GWAS. Each dataset is shown to the right. Colored panels marked with an X indicate that a given transcript is rhythmic in that dataset. B. Graph indicating the percent of rhythmic transcripts in each dataset that has also been identified to have SNPs in their loci in AD GWAS. Color of the circles indicates the P value from a Fisher’s test of the overrepresentation of AD GWAS genes present in each rhythmic dataset.

**Fig. S6:**
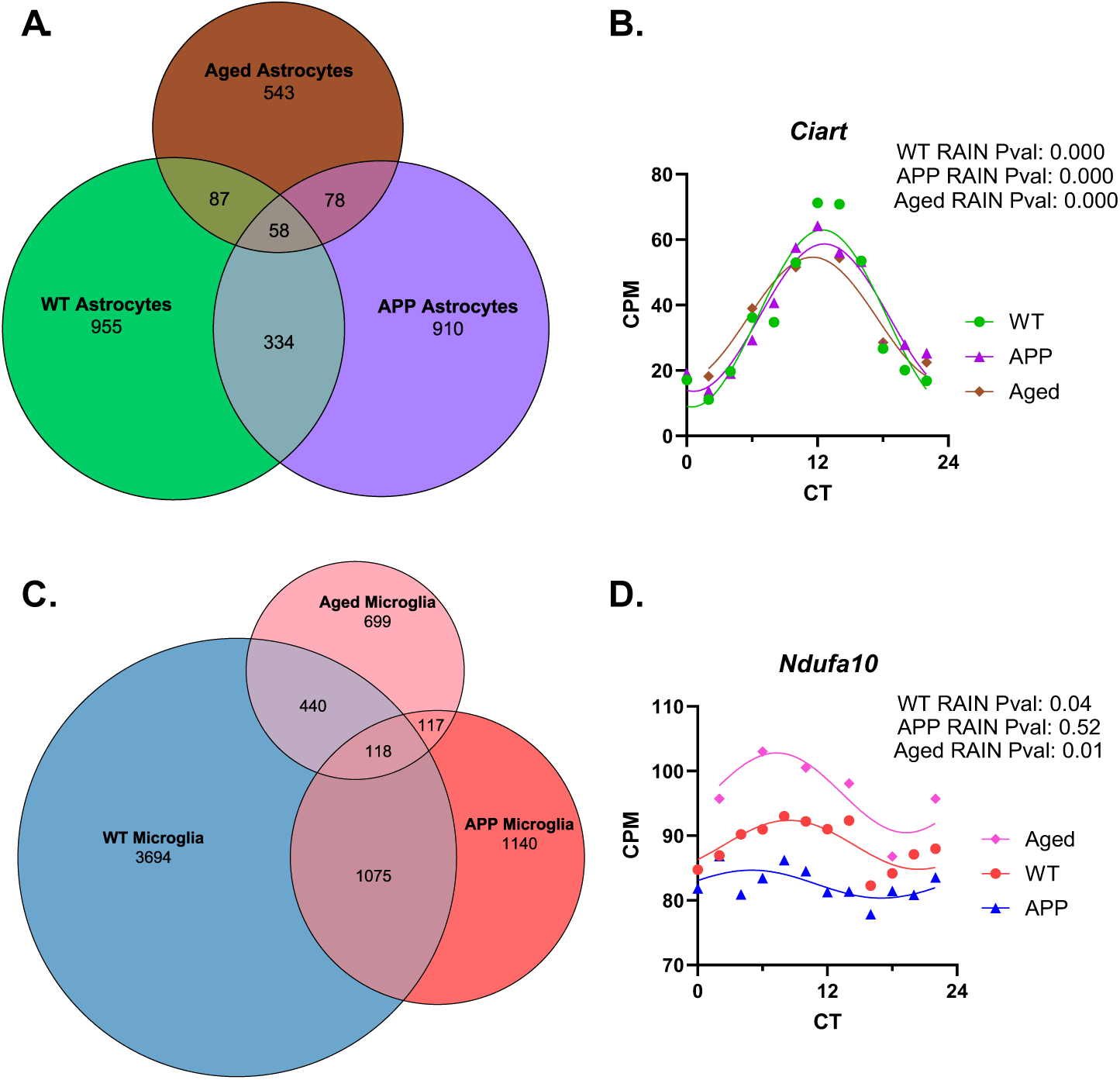
Comparison of rhythmic gene expression across disease states in astrocytes and microglia. A. Venn diagram showing minimal overlap between rhythmic genes in astrocytes from young WT, APP/PS1, and aged mice. B. Rhythmic expression of circadian gene *Ciart* is unchanged across astrocyte datasets. C. Venn diagram showing minimal overlap between rhythmic genes in microglia from young WT, APP/PS1, and aged mice. D. Rhythmic expression of electron transport chain component *Ndufa10* across microglia datasets.

**Fig. S7:**
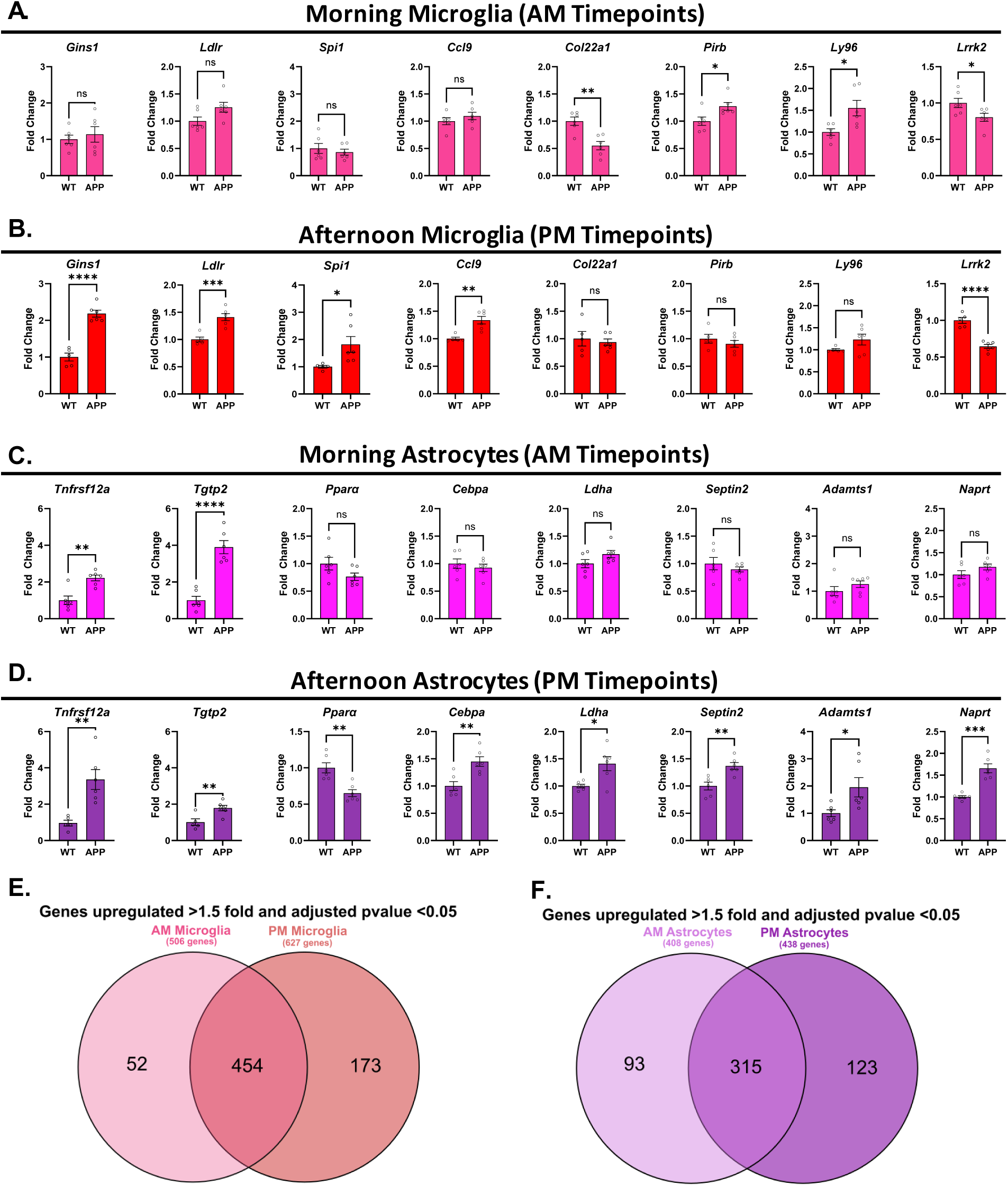
Time-of-day influences differential gene expression in APP/PS1 mice in both astrocytes and microglia. A & B. A number of representative genes plotted from microglia in the AM time bin (A) or PM time bin (B). C & D. A number of representative genes plotted from astrocytes in the AM time bin (A) or PM time bin (B). E. Venn diagram analysis of the genes with an adjusted P value <0.05 and fold change >1.5 comparing WT and APP/PS1 mice separated by the AM and PM microglia time bins. F. Venn diagram analysis of the genes with an adjusted P value <0.05 and fold change >1.5 comparing WT and APP/PS1 mice separated by the AM and PM astrocyte time bins. Graphs in (A,B,C and D) show mean ± SEM. **p < 0.01, ***p < 0.001 by unpaired two-tailed t test

**Fig. S8:**
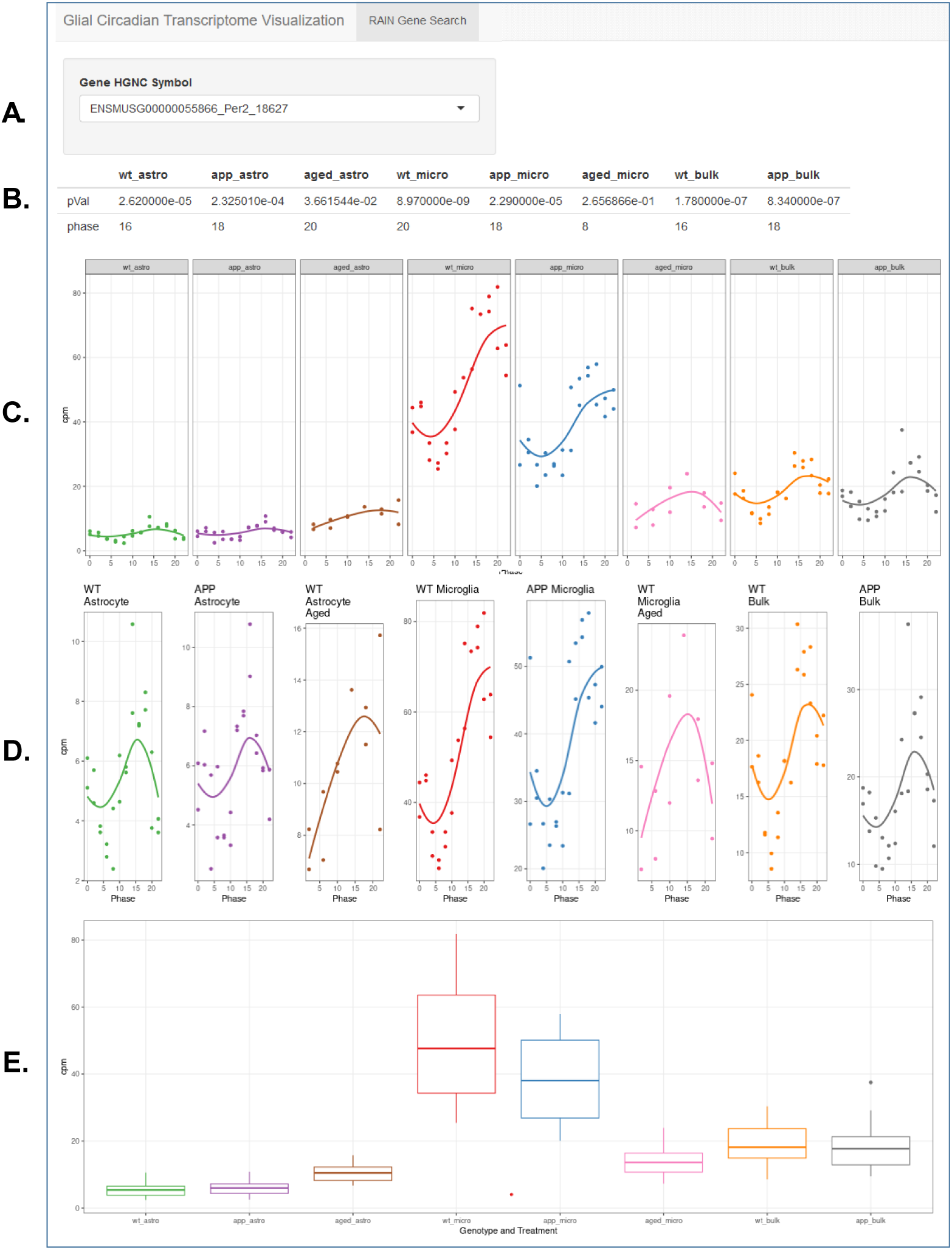
Output from Glial Circadian Transcriptome Visualization website. Typical output from https://musieklab.shinyapps.io/Glial_Circadian_Translatome/ after entry of “Per2” as gene of interest into the search field (A). RAIN-derived adjusted P values are shown along with acrophase information for each dataset in the upper right (B). Gene expression is graphed across all datasets with a fixed Y axis (C), and then on a Y-axis which is optimized for each dataset (D). Box plots of averaged gene expression across timepoints are then shown for each dataset (E).

